# *Nr6a1* controls axially-restricted body elongation, segmentation, patterning and lineage allocation

**DOI:** 10.1101/2022.03.21.485239

**Authors:** Yi-Cheng Chang, Siew Fen Lisa Wong, Jan Schroeder, Gabriel M. Hauswirth, Natalia A Shylo, Emma L Moore, Annita Achilleos, Victoria Garside, Jose M. Polo, Jan Manent, Paul Trainor, Edwina McGlinn

## Abstract

The vertebrate main-body axis is laid down during embryonic stages in an anterior-to-posterior (head-to-tail) direction, driven and supplied by posteriorly located progenitors. For the vertebral column, the process of axial progenitor cell expansion that drives elongation, and the process of segmentation which allocates progenitor-descendants into repeating pre-vertebral units, occurs seemingly uninterrupted from the first to the last vertebra. Nonetheless, there is clear developmental and evolutionary support for two discrete modules controlling processes within different axial regions: a trunk and a tail module. Here, we identify Nuclear receptor subfamily 6 group A member 1 (Nr6a1) as a master regulator of elongation, segmentation, patterning and lineage allocation specifically within the trunk region of the mouse. Both gain- and loss-of-function *in vivo* analyses revealed that the precise level of Nr6a1 acts as a rheostat, expanding or contracting vertebral number of the trunk region autonomously from other axial regions. Moreover, Nr6a1 was found to be required for segmentation, but only for trunk-forming somites, with the timely clearance of Nr6a1 critical in supporting tail formation. In parallel with these morphological outcomes, we reveal Nr6a1 as a novel regulator of global *Hox* signatures within axial progenitors, preventing the precocious expression of multiple posterior *Hox* genes as the trunk is being laid down and thus reinforcing that patterning and elongation are coordinated. Finally, our data supports a crucial role for Nr6a1 in regulating gene regulatory networks that guide cell lineage choice of axial progenitors between neural and mesodermal fate. Collectively, these data reveal an axially-restricted role for Nr6a1 in all major cellular and tissue-level events required for vertebral column formation, supporting the view that modulation of Nr6a1 expression level or function is likely to underpin evolutionary changes in axial formulae that exclusively alter the trunk region.

## Introduction

Vertebrate animals exhibit great diversity in their axial formulae, that is, the number and identity of elements that constitute the vertebral column. Axial formulae is established at early developmental stages when regional identity is superimposed on the continual process of somitogenesis (Bénazéraf and Pourquié, 2013). During primary body formation, presacral vertebrae arise from dual-fated progenitor cells known as neuro-mesodermal progenitors (NMPs) that reside in the epiblast (Cambray and Wilson, 2007; Forlani et al., 2003), with the process of gastrulation providing a constant supply of descendant cells directly to the posterior presomitic mesoderm (PSM). The resulting PSM expansion is balanced by the periodic budding of tissue from the anterior PSM, generating vertebrae precursors known as somites. As gastrulation terminates, a subset of NMPs relocate internally to the chordoneural hinge of the tailbud, where they continue to supply descendant cells destined to form post-sacral vertebrae during secondary body formation (Cambray and Wilson, 2002; Dias et al., 2020). This continues until progenitor exhaustion/PSM decline halts elongation and defines total vertebral number for that species (Denans et al., 2015; Gomez et al., 2008; Wymeersch et al., 2016). While each newly formed somite appears seemingly identical along the entire anterior-to-posterior (A-P) axis, the positional information that instructs regionally-appropriate vertebra morphology is already present prior to somite scission (Carapuço et al., 2005; Kieny et al., 1972). Central orchestrators of this positional information are the homeodomain-containing Hox proteins, a series of 39 transcription factors (in mouse/human) that work within a complex spatio-temporal hierarchy to pattern the vertebrae (Wellik, 2007). Specifically, it is the temporally-ordered activation of *Hox* gene expression within axial progenitors (Burke et al., 1995), reinforced by their genomic clustering at 4 distinct loci (Soshnikova and Duboule, 2009), that results in a spatially overlapping “Hox-code” along the A-P axis that governs regional identity (Wellik, 2007).

Evolutionary changes in vertebral number that selectively alter either the trunk or the tail/caudal regions have been repeatedly observed across the vertebrates (Buchholtz, 2007; Narita and Kuratani, 2005; Polly et al., 2001; Ward and Brainerd, 2007), supporting the view that axial elongation is driven by two consecutive developmental modules that can be manipulated independently. At a cellular level, the trunk-to-tail (T-to-T) transition is marked by a switch in axial progenitor source from epiblast- to tailbud-derived (Guillot et al., 2021; Wymeersch et al., 2016), and is coincident with a shift from an expanding to a depleting PSM size (Gomez et al., 2008). Transplantation experiments have shown that these T-to-T changes in cellular state and activity are not irreversible (Cambray and Wilson, 2002), demonstrating that intrinsic exhaustion of axial progenitors does not follow a timer mechanism and that axial identity is able to be reset to a more anterior fate based on environmental cues (Cambray and Wilson, 2002; McGrew et al., 2008). These data, together with the observation of a temporally changing transcriptomic signature of axial progenitors (Gouti et al., 2017; Wymeersch et al., 2019), as opposed to a stable signature, suggests an evolving developmental program as the main-body axis is laid down but one that can be broadly delineated into trunk and tail modules.

Gdf11 is one of the most prominent factors known to control the correct timing of the T-to-T transition. Genetic deletion of *Gdf11* (McPherron et al., 1999) or it’s receptors *Acvr2a* and *Acvr2b* (Lee et al., 2010; Oh et al., 2002) in the mouse significantly expands the trunk region, while constitutive activation of this signalling pathway had the opposite effect (Jurberg et al., 2013). However, Gdf11 signalling has pleiotropic effects across the vertebral column, with patterning changes as far anterior as the cervical region as well as tail truncation observed in *Gdf11^-/-^* mice (McPherron et al., 1999). It is currently unclear whether the cell-intrinsic mechanisms downstream of Gdf11 signalling are common across all phenotypes, though axially-restricted effectors such as Pou5f1/Oct4 in the trunk (Aires et al., 2016), and the Lin28-*let7* axis in the tail (Aires et al., 2019; Miyazawa et al., 2019; Robinton et al., 2019) have been identified.

Nuclear receptor subfamily 6 group A member 1 (Nr6a1; previously called GCNF) is an orphan-nuclear receptor whose genetic deletion in the mouse results in early embryonic lethality (Chung et al., 2001). *Nr6a1* is expressed widely within the early mouse embryo (Chung et al., 2001; Süsens et al., 1997), and is one of a select suite of marker genes that exhibits temporally-restricted expression within NMPs at embryonic day (E)8.5 (Gouti et al., 2017). Of particular note, an activating single nucleotide polymorphism (SNP) within *Nr6a1* has been associated with increased trunk vertebral number in the domesticated pig (Burgos et al., 2015; Mikawa et al., 2007; Rubin et al., 2012; Yang et al., 2009), a trait that has been selected for increased meat production (King and Roberts, 1960), with similar SNPs now found in domesticated sheep (Zhang et al., 2019). In mouse and other vertebrate species, the *Nr6a1* 3’-untranslated region harbours multiple binding sites for miR-196 and let-7 <u>(Wong et al. 2015;</u> <u>Gurtan et al. 2013; Agarwal et al. 2015)</u>, microRNA families known to constrain trunk (Wong et al., 2015) and tail (Robinton et al., 2019) vertebral number respectively, suggesting the potential for Nr6a1 to function during vertebral column formation in an axially-restricted manner, though to date, this remains to be tested.

Here, we identify Nr6a1 as a master regulator of trunk elongation, segmentation, patterning and lineage allocation in the mouse. *Nr6a1* expression within axial progenitors is dynamic, being positively reinforced by Wnt signalling at early stages and sharply terminated by the combined actions of Gdf11 and miR-196 at the T-to-T transition. Nr6a1 acts in a dose-dependent manner to control the number of thoraco-lumbar elements in a meristic not homeotic manner, with the subsequent termination of *Nr6a1* expression essential for tail development. Furthermore, Nr6a1 controls the timely progression of *Hox* expression derived from all Hox clusters. Specifically, Nr6a1 activity enhances the expression of several trunk *Hox* genes whilst temporally regulating the expression of posterior *Hox* genes. Collectively, our data reinforce the view that axial elongation is controlled by two developmental modules, the first being Nr6a1-dependent and the second Nr6a1-independent, with Nr6a1 providing central cross-talk between elongation and patterning.

## Results

### Expression and regulation of *Nr6a1* correlates with trunk formation in the mouse

The genomic locus encompassing *Nr6a1* produces multiple transcripts in the mouse (Figure S1A). These include two largely overlapping *Nr6a1* sense transcripts that produce proteins with identical DNA-binding and ligand-binding domains (Figure S1B), with a common 3’untranslated region harbouring multiple microRNA (miRNA) binding sites (Figure SlC). Within intron 3 of the long isoform, multiple antisense transcripts were identified including a long non-coding RNA (*Nr6a1os*) and a polycistronic transcript that encodes the *miR-181a2* and *miR-181b-2* microRNAs (miRNAs). Whole mount *in situ* hybridisation utilizing a riboprobe that bound both protein-coding sense transcripts revealed widespread *Nr6a1* expression at embryonic day (E) 8.5 (Figure 1A), including the posterior growth zone which houses various progenitor populations required for axial elongation (Wymeersch et al., 2016). Interrogation of a published single cell RNA-seq dataset produced from E6.5 and E8.5 embryos (Pijuan-Sala et al., 2019) confirmed robust expression of *Nr6a1* at these early stages in NMPs, caudal mesoderm and the caudolateral epiblast (Figure 1B). At E9.5, widespread *Nr6a1* expression was largely maintained consistent with previous studies (Chung et al., 2006; Fuhrmann et al., 2001), though with notable absence of expression in the heart and a visible clearing of expression from the posterior growth zone during the period known as the T-to-T transition (Figure 1A). This temporal decline in *Nr6a1* within E9.5 NMPs is supported at the single-cell level (Gouti et al., 2017). At E10.5, restricted expression became apparent within the mid- and hind- brain, cranial neural crest cells, otic vesicle and dorsal root ganglia of the trunk, with a complete absence of expression caudal to the last-formed somite. By E12.5, *Nr6a1* expression was barely detectable throughout the embryo, consistent with a “mid-gestation” pattern of expression (Gurtan et al., 2013).

**Figure 1.**
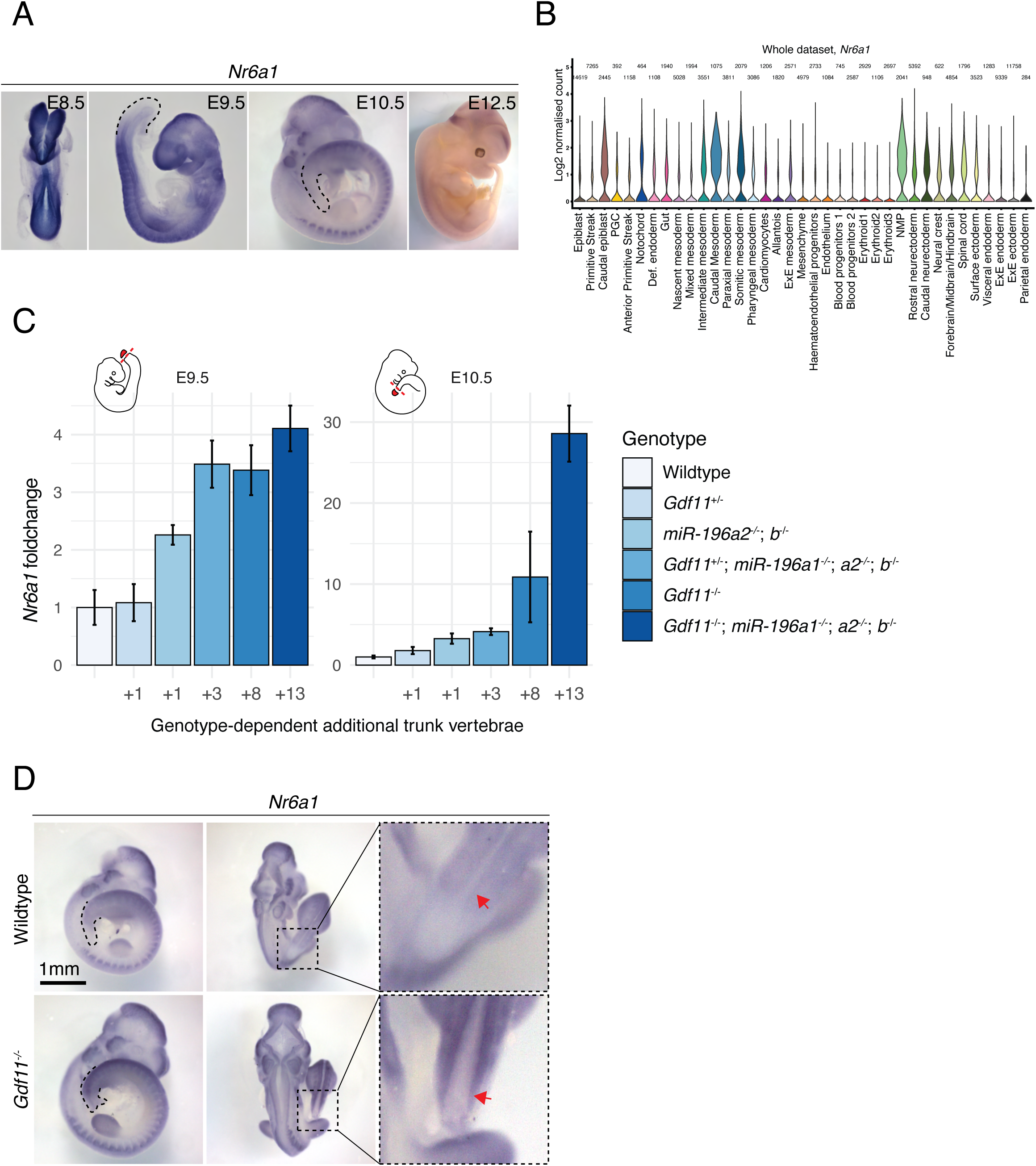
*Nr6a1* expression level positively correlates with the trunk vertebral number and is regulated by known mediators of the trunk-to-tail transition. (A) *Nr6a1* is expressed broadly at embryonics day (E)8.5, with expression becoming progressively excluded from the posterior growth zone and the heart, starting E9.5. By E10.5 *Nr6a1* exhibits tissue-specific expression, followed by complete tissue clearance by E12.5. (B) Violin plot illustrates the Log_2_ normalised count, which was derived from the raw counts from each cells divided by their size factors, of *Nr6a1* in cells of the E6.5 to E8.5 mouse embryo, revealing robust expression within neuromesodermal progenitors (NMP), caudal mesoderm and caudolateral epiblast. Data derived from (Pijuan-Sala et al., 2019). (C) Expression level of *Nr6a1* within the tailbud positively correlates with the number of trunk vertebrae that form later in development. Quantitative PCR analysis of *Nr6a1* within tailbud tissue isolated from *Gdf11* and *miR-196* single and compound mutant embryos at E9.5 and E10.5, represented as a fold change compared to wildtype (defined as 1). Sample size per genotypes ranged from 2 to 4, error bar represents standard deviation. The number of additional thoraco-lumbar vertebrae observed across the allelic deletion series (Hauswirth et al., manuscript attached) is indicated. (D) Whole mount *in situ* hybridisation reveals that the quantitative increase of *Nr6a1* in E10.5 *Gdf11^-/-^* tailbud tissue reflects a lack of timely clearance from the posterior growth zone. The arrow indicates the neural tube.

The rapid clearance of *Nr6a1* from the tailbud around E9.5 was striking, and suggested the posterior limit of its expression may be controlled by factors that position the T-to-T transition. Indeed, *Nr6a1* expression levels are known to be moderately repressed *in vivo* by *miR-196*, a factor that constrains thoraco-lumbar (T-L) number by 1 (Wong et al., 2015). To dissect *Nr6a1* regulatory mechanisms at this posterior boundary further, we took advantage of a series of recently generated mouse mutant lines which harbour an increasing number of thoraco-lumbar vertebrae dependent on the individual and combinatorial deletion of *miR-196* and *Gdf11* alleles (Hauswirth et al., manuscript attached). Quantification of *Nr6a1* expression within E9.5 tailbud tissue from across this allelic series revealed an overall positive correlation between early *Nr6a1* expression level and the number of additional T-L vertebrae that form later in development (Figure 1C). This increase in *Nr6a1* expression relative to wildtype persisted at E10.5 and in extreme genotypes, where 8 or 13 additional T-L elements ultimately form, this differential expression escalated even further. Spatial analysis of *Nr6a1* in E10.5 *Gdf11*^-/-^ embryos confirmed persistent expression in the caudal neural tube and mesoderm expanding toward the embryonic tip (Figure 1D), reminiscent of ectopic *Lin28a* and *Lin28b* expression observed in embryos of the same genotype (Aires et al., 2019). Collectively, these data revealed a dynamic pattern of *Nr6a1* expression that correlates with axial progenitors of the trunk region and one that is terminated by the synergistic action of *miR-196* and Gdf11 signaling.

### Nr6a1 is required to support trunk elongation and inhibit sacrocaudal identity

To provide insights into the mechanistic role of Nr6a1 in axial elongation, we first performed microarray transcriptomic analyses on E9.0-9.5 wildtype and *Nr6a1^-/-^* mutant embryos. The top 50 genes downregulated in *Nr6a1^-/-^* mutants compared to wildtype were analysed by ToppGene, which predicted that regionalisation, pattern specification, skeletal system development and morphogenesis, and anterior-posterior patterning were key biological processes disrupted in *Nr6a1^-/-^* mutants (Table S1 and Figure S2A). ToppGene analysis also predicted multiple phenotypes associated with abnormal morphology of the axial skeleton as likely outcomes resulting from Nr6a1 loss-of-function (Table S1). Lastly, DAVID analysis revealed that the protein families over-represented in the top 50 downregulated genes in *Nr6a1^-/-^* mutants included Homeobox proteins and Homeodomain-like proteins (Table S1) suggesting a putative link between Nr6a1 and Hox-dependent patterning mechanisms.

The early lethality of *Nr6a1*^-/-^ embryos complicates evaluation of its role during vertebral column formation (Chung et al., 2001; Lan et al., 2002). To circumvent this, we next generated a ubiquitous but temporally-controlled conditional knockout model, crossing the *Nr6a1^flx/flx^* line (Lan et al., 2003) with the *CMV-CreER*^T2^ deleter line (Sladitschek and Neveu, 2015). Low dose Tamoxifen (Tam) was administered on a single day of development from E6.25-E9.25 and skeletal analysis performed at E18.5, revealing a reduction in the number of thoracic elements in *CMV-CreER*^T2^;*Nr6a1^flx/flx^* embryos when Tam was administered at or before E8.25 (Figure S2B-C). With the time-window of action defined, we next deleted *Nr6a1* specifically within axial progenitors using the *TCreER*^T2^ deleter line (Anderson et al., 2013). Following administration of Tam at E7.5, a dose-dependent reduction in total vertebral number (TVN) and widespread vertebral patterning changes could be observed (Figure 2B and S3A). Compared to *TCreER*^T2^-positive control embryos, heterozygous conditional deletion of *Nr6a1* resulted in 2 less T elements (Figure 2B, 2D) and an overall reduction in TVN by 1 (Figure S3A) which we suggest stems from serial posteriorising homeotic transformations from the lower thoracic region onwards. For example, while the normal complement of 7 sternal rib attachments were unchanged in *TCreER*^T2^;*Nr6a1^+/flx^* embryos, the transitional vertebra (vertebra 17 (T10) in controls) was anteriorly displaced by one and all elements from the first lumbar vertebra onwards were anteriorly displaced by two. This phenotype was enhanced following homozygous conditional deletion of *Nr6a1*, with the number of sternal rib attachments reduced to 5, an overall loss of 4 T elements and a reduction in TVN by 3 (Figure 2B, 2D and S4). In these *TCreER*^T2^;*Nr6a1^flx/flx^* embryos, the positioning of rudimentary pelvic bones appeared normal relative to the last formed thoracic element, however, all intervening vertebral elements had taken on a sacral identity based on the presence of winged transverse processes, indicating a striking loss of all lumbar identity (Figure 2C). This latter result demonstrated an essential and dose-dependent role for Nr6a1 in the appropriate timing of patterning events during axial elongation, with the precocious activation of more posterior programs following Nr6a1 depletion predicted to temporally advance termination of elongation (Aires et al., 2019; Young et al., 2009) (Hauswirth et al., manuscript attached), leading to the reduction in TVN observed.

**Figure 2.**
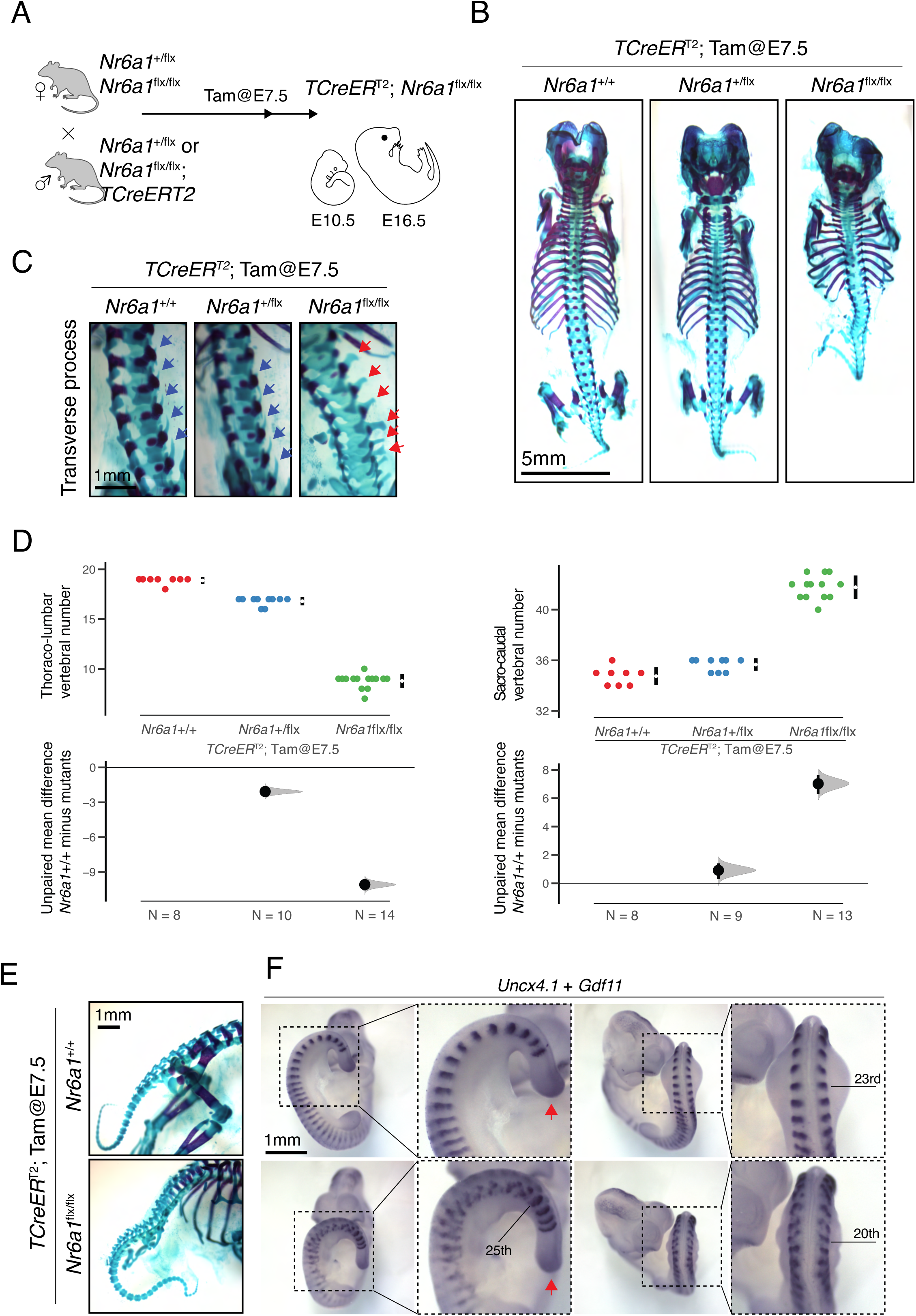
Nr6a1 is required for trunk elongation and inhibits sacro-caudal identity. (A) Conditional deletion strategy employed to remove Nr6a1 activity from within axial progenitors. Developmental stage of Tamoxifen (TAM) administration indicated. (B) Skeletal preparation of E18.5 embryos following conditional deletion of *Nr6a1* revealed a dose-dependent reduction in the number of thoraco-lumbar elements. (C) Higher magnification view of the lumbar region in embryos from (B) revealed the characteristic transverse process morphology seen in wildtype and *TCreER*^T2^; *Nr6a1^+/flx^* embryos (red arrows) had transformed to a sacral morphology in *TCreER*^T2^; *Nr6a1^flx/flx^* embryos (green arrows). (D) Quantification of thoraco-lumbar and sacro-caudal vertebral number following conditional deletion of *Nr6a1* alleles. Raw data is presented in the upper plots. Mean differences relative to wildtype (+/+) are presented in the lower plots as bootstrap sampling distributions. The mean difference for each genotype is depicted as a black dot and 95% confidence interval is indicated by the ends of the vertical error bar. (E) Analysis of caudal elements in wildtype and *TCreER*^T2^;*Nr6a1^flx/flx^* embryos revealed no difference in vertebrae morphology or number. (F) Whole mount *in situ* hybridisation for the somite marker *Uncx4.1* revealed a rostral shift in hindlimb positioning in *TCreER*^T2^;*Nr6a1^flx/flx^* embryos compared to control. Additionally, somite morphology was specifically disrupted in the future thoraco-lumbar region of *TCreER*^T2^;*Nr6a1^flx/flx^* embryos, returning to normal immediately after the hindlimb bud. Co-expression analysis of *Gdf11* (red arrows) revealed no change in expression in the tailbud between genotypes.

In addition to these patterning and numerical changes, homozygous conditional deletion of Nr6a1 resulted in rib fusion defects and vertebral malformations throughout the entire thoracic and extended-sacral regions (Figure 2B and S4). Surprisingly, however, vertebral morphology immediately after the hindlimb reverted back to normal, with tail elements of *TCreER*^T2^;*Nr6a1^flx/flx^* embryos indistinguishable from controls (Figure 2E). Consistent with this, characterisation of the early somite marker *Uncx4.1* at E10.5 revealed highly dysmorphic somite formation in the interlimb and hindlimb regions of *TCreER*^T2^;*Nr6a1^flx/flx^* embryos compared to controls, with a sharp switch back to normal appearance immediately caudal to the hindlimb bud (Figure 2F). Collectively, these striking results demonstrate an axially-restricted function for Nr6a1 in terms of vertebral number, patterning and morphology, reinforcing the view that two distinct programs govern the trunk and the tail regions - the former being Nr6a1-dependent and the latter Nr6a1-independent.

### Nr6a1 inhibits posterior *Hox* gene expression in tailbud

The sequential activation of *Hox* genes in time, and subsequently in space, underpins body patterning along the vertebrate A-P axis (Burke et al., 1995; McIntyre et al., 2007; Wellik and Capecchi, 2003). To determine whether the posteriorising homeotic transformations observed following conditional deletion of *Nr6a1* resulted from precocious activation of a posterior *Hox* code, we analysed the expression of *Hox10-13* paralogs in E9.5 tailbud tissue. Indeed, the expression of several posterior *Hox* genes were significantly up-regulated in *TCreER*^T2^;*Nr6a1^flx/flx^* tailbud compared to control, including *Hoxc12, Hoxc12, Hoxb13, Hoxc13* and *Hoxd13*, with others following a similar trend (Figure 3A). For many of these genes, a dose-dependent control of *Hox* expression levels by Nr6a1 is suggested, although not significant with current sample numbers. The most highly upregulated *Hox* gene amongst this list was *Hoxb13,* increasing more than 200-fold in *TCreER*^T2^;*Nr6a1^flx/flx^* tailbuds (Figure 3A). Spatial analysis of *Hoxb13* in wildtype E9.5 embryos (Figure 3B) confirmed expression was limited to endodermal cells of the hindgut at this stage (Zeltser et al., 1996). In contrast, E9.5 *TCreER*^T2^;*Nr6a1^flx/flx^* embryos showed additional ectopic expression in the region of the tailbud where NMPs are known to reside (Wymeersch et al., 2016) (Figure 3B). Ectopic *Hoxb13* expression persisted in E10.5 *TCreER*^T2^;*Nr6a1^flx/flx^* embryos throughout the tailbud mesoderm including the NMP-containing chordoneural hinge, and increased *Hoxb13* expression levels were observed in the neural tube relative to wildtype (Figure 3B). These data reveal an essential role for Nr6a1 in ensuring correct spatio-temporal activation of a collective posterior *Hox* code within the tailbud and, specifically, in preventing their precocious expression at stages prior to the T-to-T transition.

**Figure 3.**
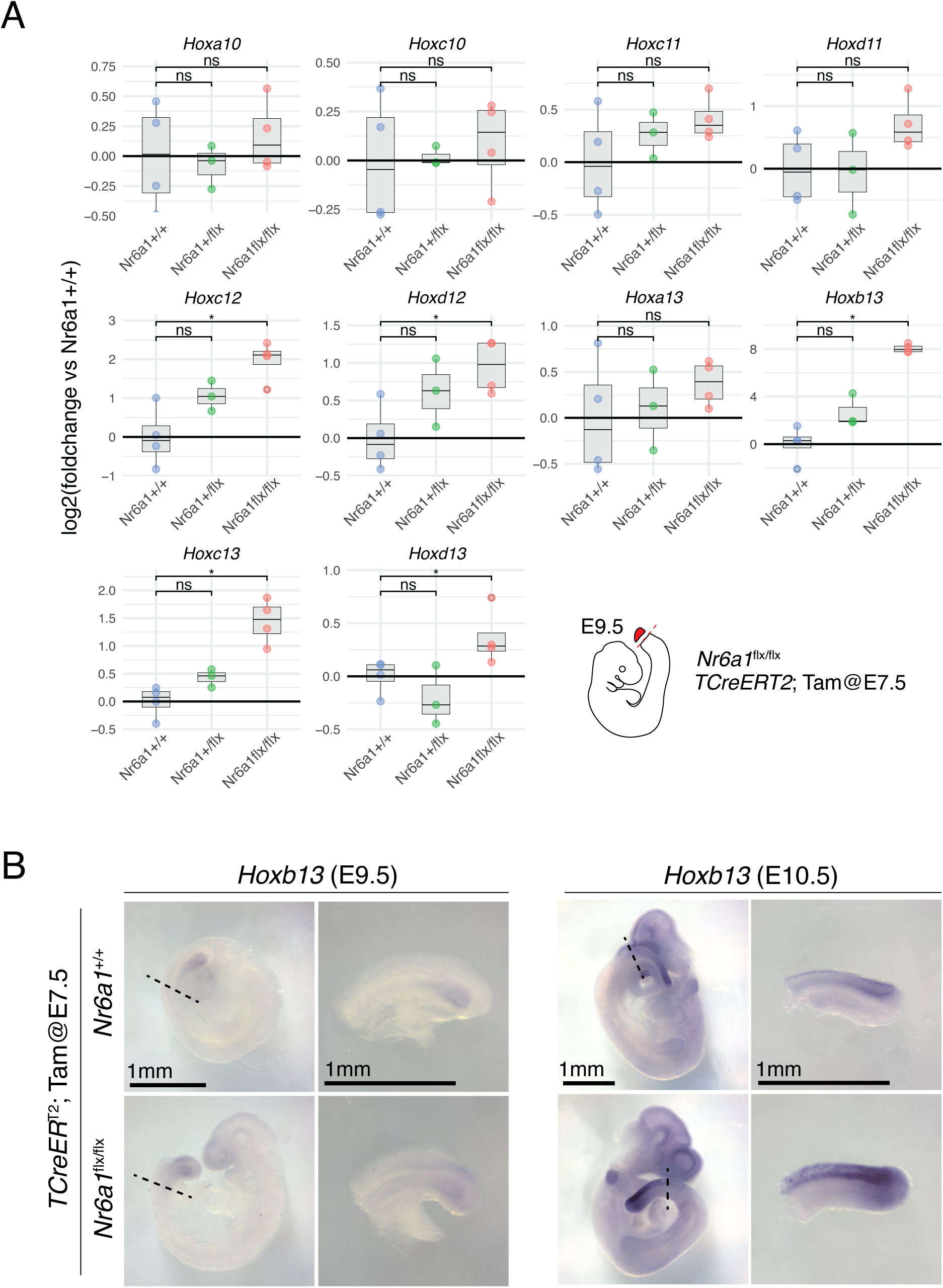
Nr6a1 suppresses posterior *Hox* gene expression. (A) Quantitative PCR analysis of posterior *Hox* gene expression within tailbud tissue isolated from *TCreER*^T2^ controls, *TCreER*^T2^;*Nr6a1*^+/flx^ and *TCreER*^T2^;*Nr6a1^f^*^lx/flx^ embryos at E9.5, n = 3/genotype. Statistical analysis performed using Wilcoxon test, * p<0.05, ** p< 0.01. (B) Whole mount *in situ* hybridisation revealed a spatial expansion of *Hoxb13* expression in *TCreER*^T2^;*Nr6a1*^+/flx^ embryos compared to wildtype, at both E9.5 and E10.5. Red arrow indicates the region containing the NMP population.

### Nr6a1 clearance is essential for the trunk-to-tail transition

During development, the clearance of expression signatures can be instructive or passive. To test the morphological and molecular importance of timely Nr6a1 clearance at the T-to-T transition, we next turned to a gain-of-function approach where transgenic expression of *Nr6a1* downstream of *Cdx2* regulatory elements (Benahmed et al., 2008) maintained gene expression in the tailbud throughout axial elongation (*Cdx2P:Nr6a1*). Skeletal analysis of *Cdx2P:Nr6a1* embryos at E18.5 revealed thoraco-lumbar expansion, constituting an additional 2 T and 2-4 L vertebrae relative to the wildtypes (Figure 4A). As the number of sternal rib attachments was unaltered in these transgenics (Figure S5A), the additional T elements are considered to be of caudal thoracic identity. Within the *Cdx2P:Nr6a1* extended lumbar region, almost all of the elements harboured an accessory process known as the anapophysis, a feature usually restricted to the first 3 lumbar elements in wildtype (Figure 4B). As such, we can pinpoint the presacral expansion in *Cdx2P:Nr6a1* embryos to elements immediately surrounding the T-L junction, in terms of their identity. Increased Nr6a1 levels did not overtly affect vertebrae morphology within the T-L region up until the most caudal lumbar elements, after which, all sacro-caudal elements became small, fused and exhibited defects in dorsal closure (Figure 4A and S5B). These regionally-restricted phenotypic changes were complementary to those observed following conditional deletion of Nr6a1 (Figure 2B), and demonstrate that the normal clearance of Nr6a1 at the T-to-T transition is not an inconsequential outcome of an altered developmental program but rather, Nr6a1 clearance is essential for correct sacral positioning and normal tail morphology.

**Figure 4.**
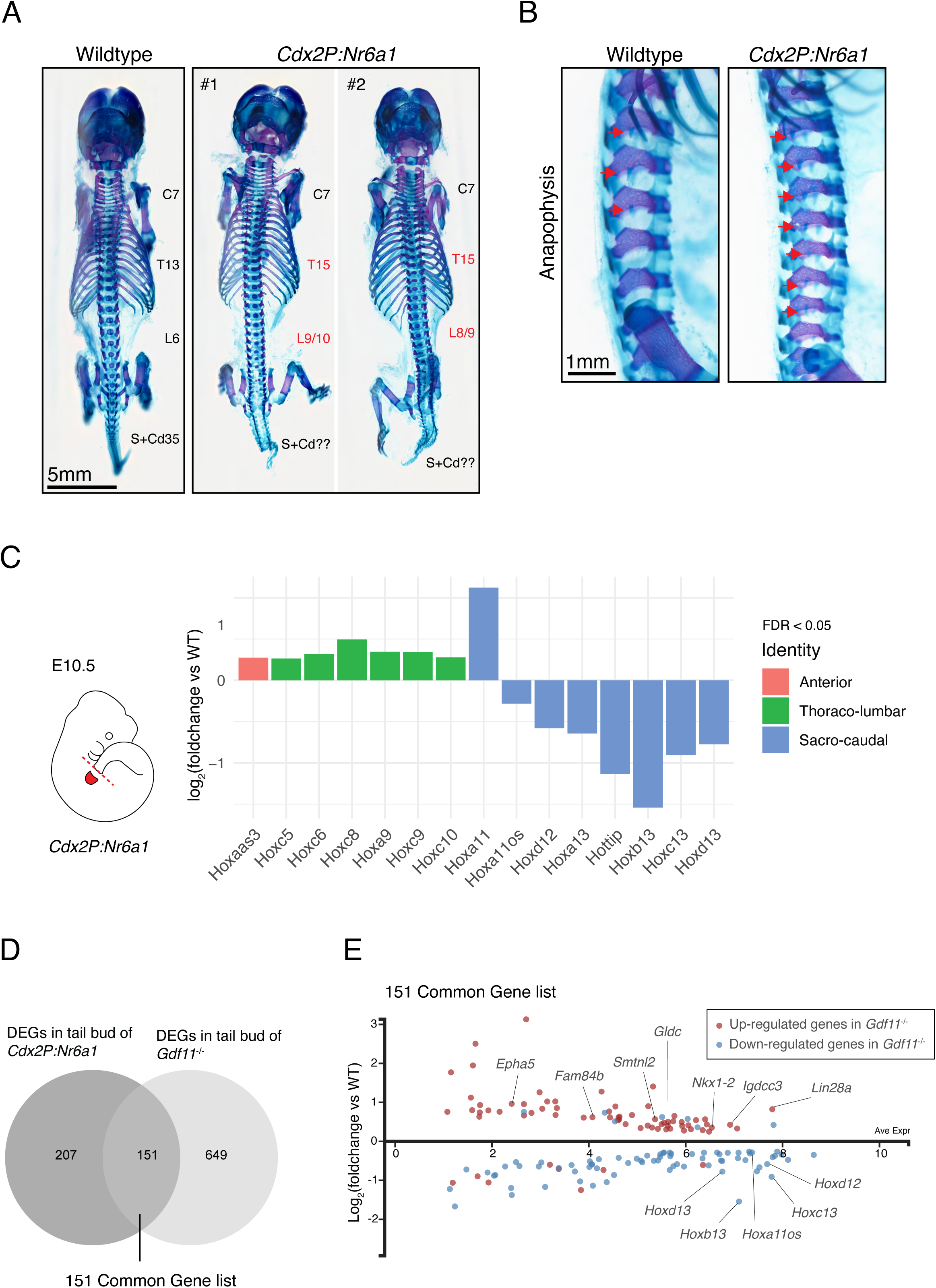
Prolonged Nr6a1 activity is sufficient to increase thoraco-lumbar count and sustain trunk expression signatures. (A) Skeletal preparation of E18.5 embryos, dorsal view, revealed an increase in the number of thoraco-lumbar elements in *Cdx2P:Nr6a1* embryos compared to wildtype; C = cervical, T = thoracic, L = lumbar, S = sacral and Cd = caudal vertebrae. (B) Higher magnification, lateral view, of the lumbar region in embryos from (A) revealed the anapophysis (red arrow) normally present on the first 3 lumbar elements in wildtype was now present on 7 of the 8 lumbar elements in *Cdx2P:Nr6a1* embryos. (C) RNAseq analysis of E10.5 wildtype and *Cdx2P:Nr6a1* tailbuds revealed widespread changes in *Hox* expression downstream of Nr6a1. Results are presented as a log2- transformed fold change in *Cdx2P:Nr6a1* samples relative to wildtype, n=2 for *Cdx2P:Nr6a1* and n=4 for wildtype. Only those *Hox* genes with a false discovery rate (FDR) < 0.05 are displayed, and are colour-coded based on the axial region where the Hox protein functions. (D) Venn diagram overlay of differentially expressed genes (DEGs) in *Cdx2P-Nr6a1* (this work) and *Gdf11^-/-^* (Aires et al., 2019) tailbud tissue reveals substantial overlap. (E) Analysis of the 151 co-regulated genes identified in (D), presented here as log2- transformed fold change in *Cdx2P:Nr6a1* samples relative to wildtype, revealed that 136 genes (90%) displayed the same direction of regulation in both genetically-altered backgrounds. Red and blue dots indicate up-regulated and down-regulated genes in *Gdf11^-/-^* tailbuds, respectively.

To ascertain whether *Hox* expression dynamics were altered in *Cdx2P:Nr6a1* embryos, we performed RNA-seq analysis of tailbud tissue at E10.5, a time when Nr6a1 should no longer be expressed at this site. Compared to wildtype controls, the prolonged maintenance of Nr6a1 activity resulted in a modest upregulation of several trunk *Hox* genes, including *Hoxc6*, *Hoxc8*, *Hoxa9*, *Hoxc9* and *Hoxc10*, genes known to participate in conveying T-L identity (van den Akker et al., 2001; McIntyre et al., 2007; Wellik and Capecchi, 2003) (Figure 4C). Conversely, genes of the *Hox11-13* paralog groups and two 5’-Hox cluster antisense genes *Hoxa11os* and *Hottip* showed reduced expression in *Cdx2P:Nr6a1* tailbuds (Figure 4C), with *Hoxb13* again having the largest differential response to altering Nr6a1 levels. One exception to this trend was *Hoxa11*, which was found to be upregulated in *Cdx2P:Nr6a1* tailbuds. A similar inverse relationship between *Hoxa11* and other posterior *Hox* cluster genes, including *Hoxa11os*, has also been observed in E10.5 *Gdf11*;*miR-196* single and compound mutant mice which harbour additional thoraco-lumbar elements (Hauswirth et al., manuscript attached). An antagonistic relationship between *Hoxa11os* and *Hoxa11* exists within the early mouse limb bud, resolving into mutually exclusive domains of expression as the limb bud elongates (Kherdjemil et al., 2016). In this context, *Hoxa11os* transcription is dependent on Hoxa13/d13 and suppresses *Hoxa11*, with our data supporting the potential for a parallel regulatory network within the paraxial mesoderm. In summary, we show that Nr6a1 alone is sufficient to control the temporal progression of *Hox* activation of all 4 *Hox* clusters. Nr6a1 activity positively reinforces trunk identity whilst suppressing precocious activation of caudal fate during trunk elongation, keeping the two developmental modules distinct. This control over *Hox* cluster progression by Nr6a1 correlated well with altered patterning outcomes observed following manipulation of Nr6a1 levels. However, the increase in T-L vertebral number seen in *Cdx2P:Nr6a1* embryos was harder to reconcile, since meristic changes (i.e. changes in vertebral number) are historically not thought to ensue downstream of changes to a single *Hox* gene or Hox paralog group.

### Opposing effects of Nr6a1 and Gdf11 signalling on a core trunk regulatory module

The similarity of T-L expansion phenotypes seen in Nr6a1 gain-of-function (Figure 4A) and Gdf11 loss-of-function (McPherron et al., 1999) mice prompted us to compare the global molecular changes identified in our E10.5 *Cdx2P:Nr6a1* RNAseq dataset with those of a published E10.5 *Gdf11^-/-^* tailbud dataset (Aires et al., 2019). This analysis revealed 151 conserved differentially expressed genes (Figure 4D), 90% of which displayed the same direction of regulation in both mutants (Figure 4E). 63 of the 136 co-regulated genes were up-regulated, including genes enriched in NMPs (*Nkx1-2*, *Epha5, Gpm6a* and *Gldc*) *(Gouti et al., 2017)*, as well as genes known to promote axial elongation such as *Lin28a*, *Lin28b* (Aires et al., 2019; Miyazawa et al., 2019; Robinton et al., 2019). On the other hand, 73 of the 136 co-regulated genes were down-regulated and included many posterior *Hox* genes, supporting an antagonistic arrangement whereby Nr6a1 promotes and Gdf11 terminates a core trunk gene regulatory network.

To test the *in vivo* epistatic interaction between Nr6a1 and Gdf11, we set out to assess whether Nr6a1 mediates Gdf11-dependent phenotypes through compound mutant analysis. Conditional deletion of Nr6a1 using the *T-CreER*^T2^ deleter line was repeated with Tam administration at E8.5, one day later than previous experiments and at a time when deletion was anticipated to have little to no phenotypic consequence that would confound compound mutant analysis. Indeed, no change in TVN was observed in *TCreER*^T2^;*Nr6a1^flx/flx^* embryos when Tam was administered at E8.5, on an otherwise wildtype (*Gdf11^+/+^*) or *Gdf11^+/-^* background (Figure S3A). However on a *Gdf11^-/-^* background which is known to display an expanded T-L count of 27 vertebrae (T18 L9; Figure 5B-C; (McPherron et al., 1999), conditional deletion of Nr6a1 at E8.5 significantly rescued T-L count by 3-4 elements (T16/17 L7), with Nr6a1 dose-dependent effects revealed (Figure 5B-C). This temporal deletion of Nr6a1 did not rescue aberrant tail vertebrae morphology nor tail truncation known to occur in *Gdf11^-/-^* embryos, consistent with earlier interpretation that Nr6a1 function was not required at this site. This work identifies separable phenotypes, trunk elongation and tail morphology, that each depend on Gdf11 activity but utilise different downstream mechanisms. Collectively, this work establishes Nr6a1 as a master intrinsic regulator of T-L vertebral number and morphology, and reveals that termination of primary body axis formation by Gdf11 is mediated, likely to a large extent, through the clearance of *Nr6a1* expression.

**Figure 5.**
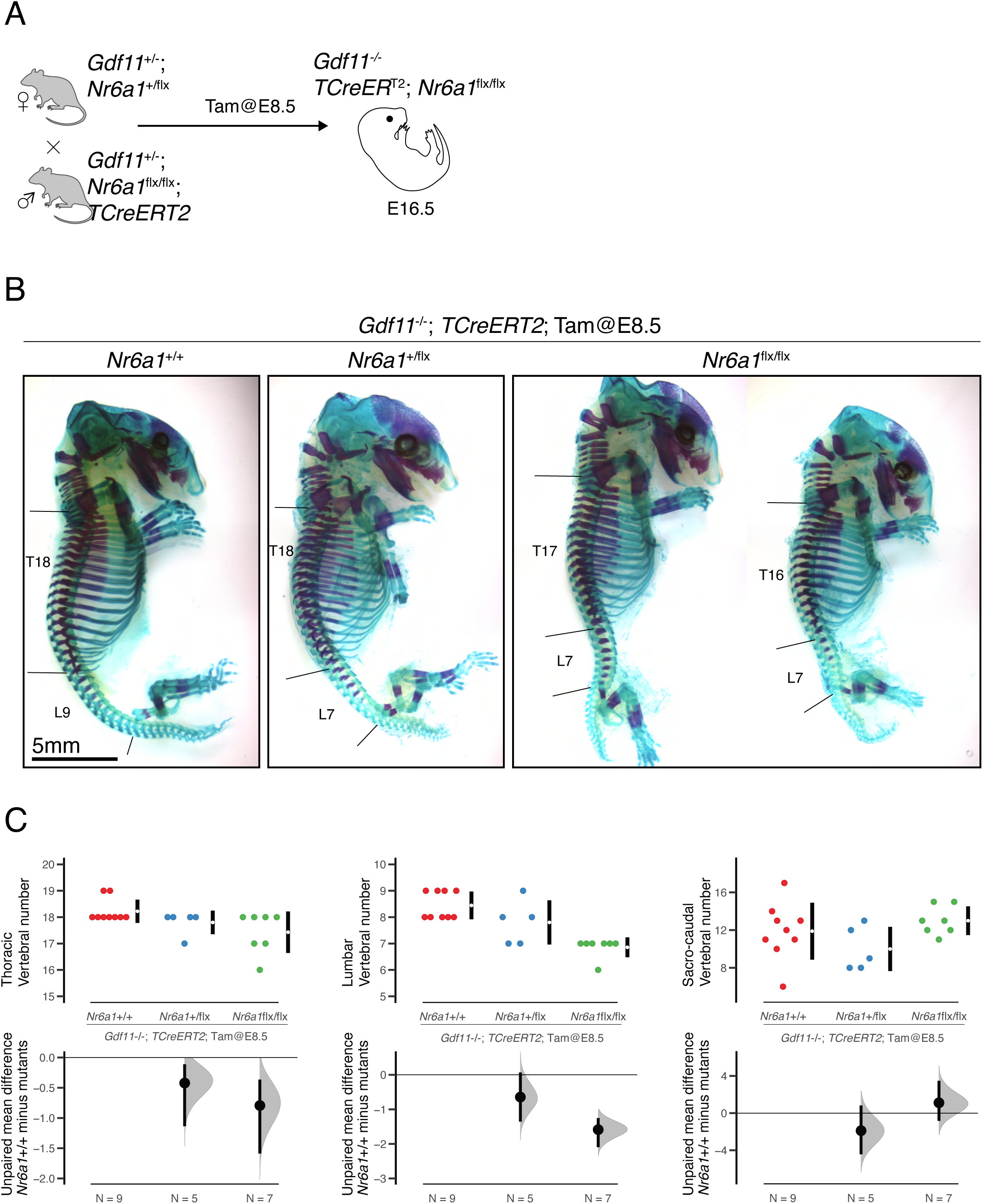
Gdf11 signalling terminates Nr6a1-dependent trunk elongation. (A) *TCreER*^T2^*;Nr6a1* conditional deletion strategy on a *Gdf11* mutant background. (B) Skeletal preparation of E18.5 embryos, lateral view, revealed a dose-dependent rescue of *Gdf11^-/-^* thoraco-lumbar expansion following conditional deletion of *Nr6a1* alleles. T = thoracic, L = lumbar. (C) Quantification of thoracic, lumbar and sacro-caudal vertebral number across genotypes. Raw data is presented in the upper plots. Mean differences were calculated relative to *Gdf11^-/-^*;*Nr6a1^+/+^* and presented in the lower plots as bootstrap sampling distributions. The mean difference for each genotype is depicted as a black dot and 95% confidence interval is indicated by the ends of the vertical error bar.

### Nr6a1, in parallel with the external factors, regulates timing of collinear *Hox* expression

To more precisely delineate the upstream signals influencing *Nr6a1* expression, and the downstream molecular events requiring Nr6a1 activity, we turned to an *in vitro* embryonic stem cell (ESC) differentiation approach that models the developmental kinetics of a posterior growth zone by the sequential addition of FGF2 on day (d) 0, WNT pathway agonist CHIR99021 on d2, with or without the addition of Gdf11 on d3 (Figure S6A; based on (Gouti et al., 2014, 2017). Induction of NMP identity and 3’ *Hox* cluster activation was confirmed soon after Wnt activation, collinear trunk *Hox* activation ensued quickly thereafter, and progression through to a posterior *Hox* code required Gdf11 (Figure S6B). In wildtype cells, *Nr6a1* was present in epiblast-like d2 cells and its expression was substantially enhanced on d2.5 and d3 following the induction of an NMP-like state (Figure S6C). *Nr6a1*, together with other signature genes of trunk-producing (E8.5) NMP (Gouti et al., 2017), declined as cells were cultured for a further 24h (d4) (Figure S6D). Conversely, signature genes of the sacro-caudal producing (E9.5) NMP increased expression on d4 (Figure S6D). While *Nr6a1* expression was already decreasing after d3, it was sharply terminated following addition of exogenous Gdf11 (Figure S6C), consistent with earlier *in vivo* results. Interestingly, an inverse temporal pattern of expression was observed for *Nr6a1os*, remaining very low until the addition of Gdf11 when levels increased 4-fold (Figure S6C). We find that the promoter of *Nr6a1os* is accessible in epiblast stem cells treated with or without Chiron for 48h (Figure S7A) (Neijts et al., 2016), and harbours several consensus binding motifs for Smad2/3 (Figure S7B), ultimate transducers of Gdf11 signalling (Oh et al., 2002) and Smad4, the common-mediator Smad, which forms heterotrimeric complex with Smad2/3 to control the target genes (Massagué et al., 2005). This raises the intriguing possibility that Gdf11 suppression of *Nr6a1* sense transcripts is in fact mediated via a direct activation of *Nr6a1os* and subsequent genomic interactions between sense and antisense loci that establish mutually exclusive expression patterns, as has been observed for other developmental loci (Kherdjemil et al., 2016).

With wildtype differentiation conditions established, we next generated *Nr6a1^-/-^* ESC clones by CRISPR/Cas9 (Arbab and Sherwood, 2016; Arbab et al., 2015), deleting the DNA-binding domain from both alleles (Figure S8). These *Nr6a1^-/-^* ESCs were able to generate NMPs with equal kinetics to wildtype cells (Figure S9), however, RNAseq analysis revealed clear transcriptomic changes as differentiation proceeded. The robust initiation of a trunk Hox code normally seen 24 hours after Wnt activation was reduced in *Nr6a1*^-/-^ cells particularly for genes of the *Hoxa, Hoxc* and *Hoxd* clusters, with the expression of *Hoxc6*, *Hoxa7*, *Hoxc8*, *Hoxa9*, *Hoxc9* and *Hoxd9* all diminished relative to WT on d3 (Figure 6A). This relative reduction was exacerbated even further when cells were cultured in Fgf/Wnt alone for a further 24 hours. Genes of the *Hoxb* cluster showed mixed responses in *Nr6a1*^-/-^ cells, however collectively, these observed expression signatures support the view that Nr6a1 does not serve a major role in the initiation of trunk Hox genes but is required to achieve their full expression potential.

**Figure 6.**
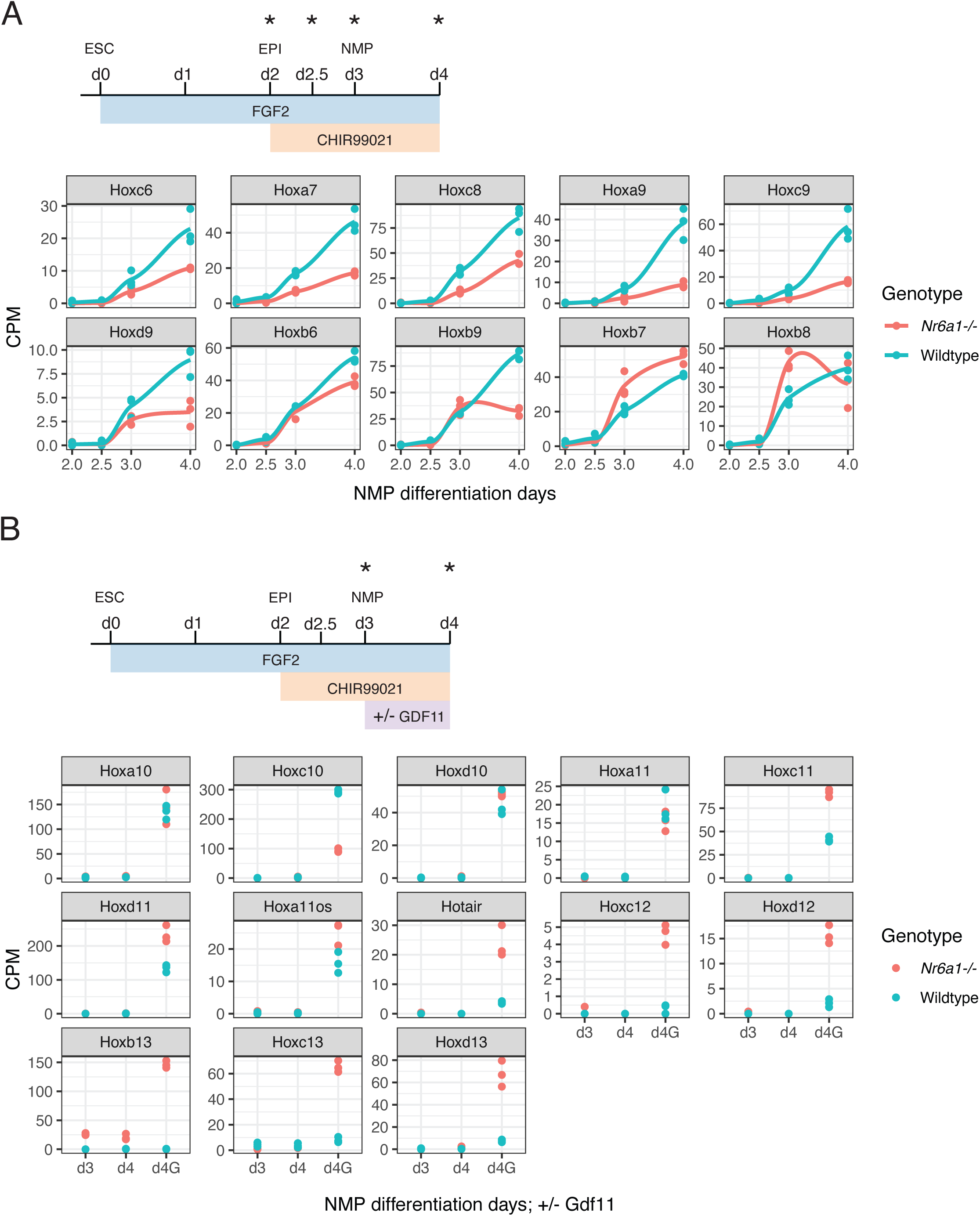
*Nr6a1* and Gdf11 signalling cooperate in the control of *Hox* cluster transitions. (A-B) Differential *Hox* expression kinetics observed during *in vitro* differentiation of *Nr6a1*^-/-^ and wildtype ESCs. (A) RNAseq analysis of trunk *Hox* gene expression (*Hox6*-*9*) in *Nr6a1^-/-^* (red) and wild type (blue) cells. *In vitro* differentiation protocol schematised, analysis timepoints for these genes marked with asterisk and data represented as counts per million (CPM). Only those genes showing a false discovery rate (FDR) < 0.05 were visualised. (B) RNAseq analysis of posterior *Hox* gene expression (*Hox10*-*13*) in *Nr6a1^-/-^* (red) and wild type (blue) cells. *In vitro* differentiation protocol schematised, analysis timepoints for these genes marked with asterisk and data represented as counts per million (CPM). Only those genes showing a false discovery rate (FDR) < 0.05 were visualised. G=addition of Gdf11.

Under these standard Fgf/Wnt culture conditions, the activation of posterior *Hox10*-*13* genes was almost universally absent in cells of both genotypes (Figure 6B). For wildtype cells this is expected, since Gdf11 is required in these cultures for robust activation (Figure S6B) (Jansz et al., 2018; Lippmann et al., 2015). For *Nr6a1*^-/-^ cells, however, these results allowed us to build a molecular hierarchy which was not able to be deciphered from *in vivo* experiments: the loss of Nr6a1 is not sufficient on its own to activate posterior *Hox* expression, and requires positive input from additional signal(s). Once Gdf11 was exogenously applied to these cultures, *Nr6a1*^-/-^ cells activated posterior *Hox* expression with far greater amplitude than wildtype cells, with this heightened response to Gdfll more evident the further 5’ within a cluster the gene was located (Figures 6B and S10). One striking exception to this identified hierarchy was *Hoxb13*. 24 hours after the activation of Wnt, and without any exogenous Gdf11, *Hoxb13* expression was induced 25-fold relative to wildtype cells where a complete absence of expression was expected and observed (Figure 6B). Expression persisted at this level without further signal input, and upon addition of Gdf11, *Hoxb13* showed the largest differential increase in expression relative to wildtype of any posterior *Hox* gene (150-fold). The mechanism behind this unique regulation of *Hoxb13* versus other posterior *Hox* genes remains to be defined, but is consistent with the heightened regulation of *Hoxb13* in Nr6a1 *in vivo* gain-of-function experiments. With this exception noted, our *in vitro* results globally reinforce the dominant role of Gdf11 in activating posterior *Hox* expression, but build in an essential function for Nr6a1 in suppressing the untimely/precocious activation of posterior *Hox* expression and in defining their precise levels once expression has been activated.

### *Nr6a1* biases cell lineage choice *in vitro* and *in vivo*

The allocation of NMP descendants towards neural or mesodermal lineages is guided by both extrinsic (Martin and Kimelman, 2012; Olivera-Martinez and Storey, 2007; Olivera-Martinez et al., 2012) and intrinsic factors (Koch et al., 2017; Takemoto et al., 2011), with a disproportionate allocation to one lineage often resulting in termination of elongation (Chapman and Papaioannou, 1998; Nowotschin et al., 2012). Progenitor cells within the *Gdf11*^-/-^ tailbud are biased toward a neural fate and an enlarged neural tube forms within caudal regions of these truncated mutant embryos (Aires et al., 2019). In other developmental contexts, Nr6a1 activity has been linked to neural specification and differentiation *in vitro* (Akamatsu et al., 2009; Gu et al., 2005) and *in vivo* during mouse (Chung et al., 2006) and *Xenopus* (Barreto et al., 2003a, 2003b) development, raising the possibility that Nr6a1 may promote neural cell fate within bipotent NMPs. To address this, we analysed the expression of genes previously found to delineate mesodermal progenitors (MP) or neural progenitors (NP) of the E8.5 tailbud (Gouti et al., 2017) within our various RNAseq datasets. Interrogation of *in vitro* differentiation samples 24 hours after Wnt activation revealed heightened expression of MP genes and reduced expression of NP genes within *Nr6a1*^-/-^ cells compared to wildtype (Figure 7A). Conversely, interrogation of *in vivo Cdx2P:Nr6a1* tail buds revealed the opposite, all NP genes were found to be up-regulated while all MP genes down-regulated when Nr6a1 was ectopically maintained (Figure 7B). Together, these results identify *Nr6a1* as a novel intrinsic factor essential for regulating the balance between neural and mesodermal fates within NMPs during axial elongation.

**Figure 7.**
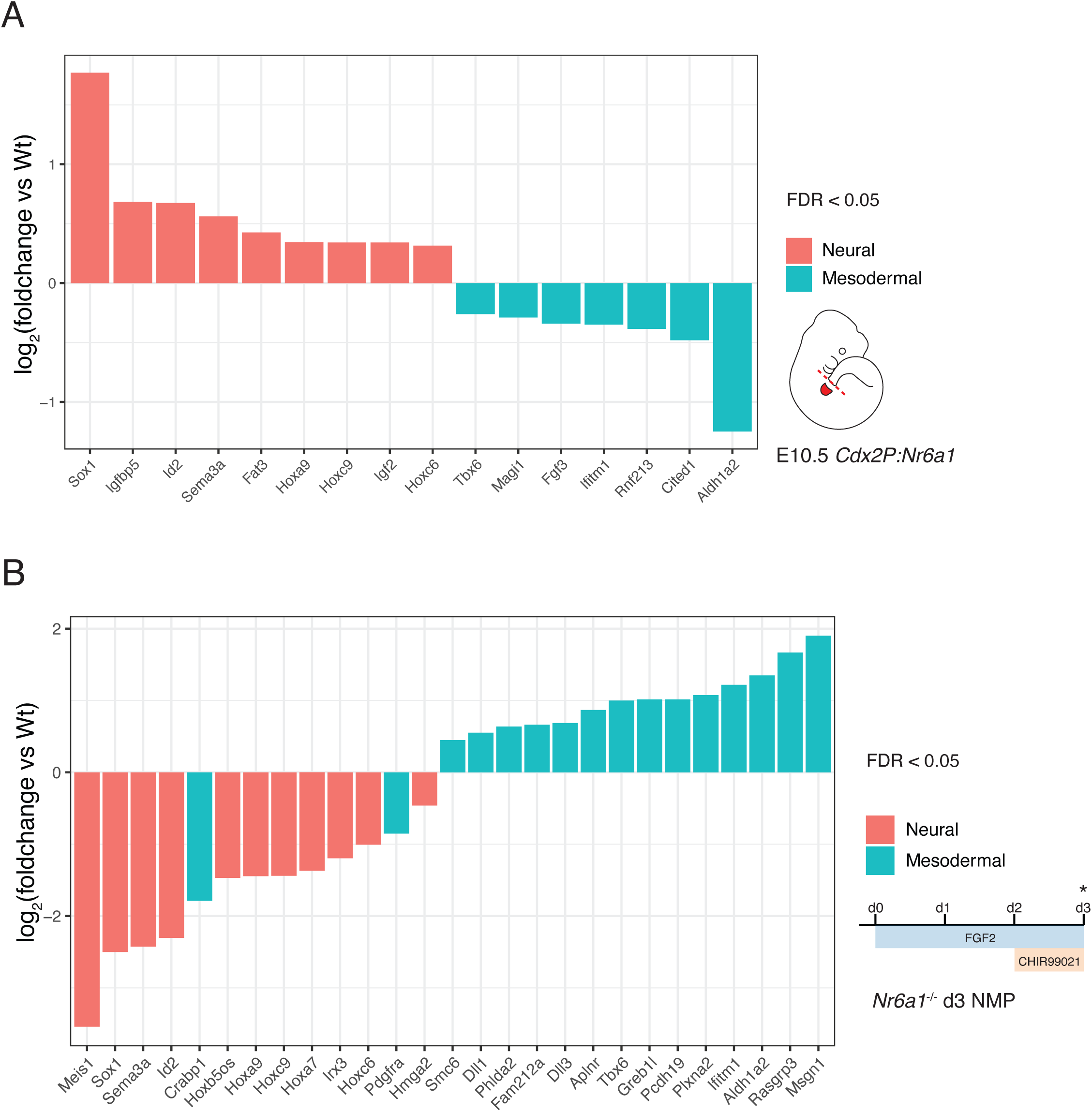
Nr6a1 activity regulates neural vs mesodermal expression signatures *in vitro* and *in vivo*. (A) Heightened neural (red) and reduced mesodermal (blue) gene expression signatures were observed following prolonged maintenance of Nr6a1 within the mouse tailbud. RNAseq analysis was presented as a log2-transformed fold change in *Cdx2P:Nr6a1* samples relative to wildtype, n=2 for *Cdx2P:Nr6a1* and n=4 for wildtype. Neural and mesodermal gene lists were selected based on *in vivo* single cell RNAseq analysis (Gouti et al., 2017), and only genes with a false discovery rate (FDR) < 0.05 are displayed. (B) Heightened mesodermal (blue) and reduced neural (red) gene expression signatures were observed in *Nr6a1^-/-^ in vitro*-derived NMPs compared to wildtype. *In vitro* differentiation protocol schematised, analysis timepoint (d3) marked with asterisk. RNAseq analysis is presented as a log2-transformed fold change in *Nr6a1^-/-^* samples relative to wildtype, n=3/genotype. Neural and mesodermal gene lists were selected based on X, and only genes with a false discovery rate (FDR) < 0.05 are displayed.

## Discussion

The initial stages of vertebral column formation require many cellular decisions and tissue-level processes that, despite their required coordination, are often considered separately in terms of genetic regulation. Here, we have identified a single factor that coordinately controls vertebral number and identity, somite segmentation and NMP cell lineage choice in the mouse. The axially-restricted manner in which Nr6a1 functions reinforces the existence of distinct developmental programs controlling trunk vs. tail formation, emphasising that this is not a developmental continuum that is delineated by a switch in patterning program and progenitor location alone. Rather, this switch involves additional changes in the genetic regulation of segmentation and regional control of vertebral number, all processes unified in their requirement for precise levels of Nr6a1 activity.

Complete loss of Nr6a1 in the mouse was found to slow the overall pace of development soon after the initiation of somitogenesis and, morphologically, these null embryos failed to progress through the period of embryonic turning (E8.75) (Chung et al., 2001). Nonetheless, up until approximately E9.5 when *Nr6a1^-/-^* embryos died, the very anterior and posterior embryonic ends continued to expand and differentiate while the trunk region stalled, consistent with our collective analyses establishing Nr6a1 as a master regulator of trunk development. *Nr6a1* transcripts have been detected ubiquitously in the mouse embryo at E6.5 (Chung et al., 2001; Fuhrmann et al., 2001), however, the first 7-13 somites still form in the complete absence of Nr6a1 activity (Chung et al., 2001), broadly correlating with the future cervico-thoracic boundary. Here, *Nr6a1* expression levels rise sharply in cells of the posterior embryo central to axial elongation (Pijuan-Sala et al., 2019), a site high in Wnt activity (Mohamed et al., 2004) which we show *in vitro* has the ability to rapidly enhance *Nr6a1* transcript levels. As the T-L region is being laid down, Nr6a1 levels remain high, and a striking correlation between caudal *Nr6a1* expression level and total T-L vertebral number was observed across our *miR-196- TKO;Gdf11^-/-^* deletion series. Indeed, *Nr6a1* levels were shown to be quantitatively instructive based on the dose-dependent phenotypes observed in both Nr6a1 gain- and loss-of-function scenarios. As development proceeds and the primary body elongation program comes to its eventual completion, we show an essential requirement for the clearance of *Nr6a1* expression within the tailbud. This clearance was achieved by multiple mechanisms, including the rising levels of Gdf11 signalling as well as microRNA repression by the posteriorly-expressed miR-196 paralogs. The 3’UTR of *Nr6a1* in fact houses binding sites for several microRNAs, including another Hox-embedded miRNA family, *miR-10*, as well as the developmental timing microRNA family, *let-7* (Figure S1C). *miR-10* paralogs are genomically positioned between *Hox4*/*5* paralogs and thus are expressed from early stages of axial elongation (Kloosterman et al., 2006; Mansfield et al., 2004). This raises the intriguing possibility that miR-10 may act to buffer the functional domain of Nr6a1 activity at more anterior locations, while miR-196 reinforces its posterior boundary, collectively delimiting Nr6a1 output to the T-L region in synchrony with the temporal progression of Hox cluster activation. Finally, within the tail-forming region, the rising levels of Gdf11, and decreasing levels of Nr6a1 activates expression of progressively more posterior *Hox* genes, including the Hox13 paralogs which shift the Lin28-*let-7* axis in favour of *let-7* expression (Aires et al., 2019), a potent repressor of Nr6a1 at least *in vitro* (Gurtan et al., 2013) thus reinforcing termination of expression. Collectively, this work has revealed a highly integrated series of regulatory mechanisms defining axially-restricted expression of *Nr6a1*, and ultimately, defining vertebral number within the trunk region of the mouse.

The consequences of sustaining Nr6a1 activity *in vivo* for longer than normal bared striking resemblance to those observed following sustained activity of the Pou-domain transcription factor Oct4/Pou5f1, both in terms of expanded trunk vertebral number and in the delayed activation of posterior *Hox* expression (Aires et al., 2016). This parallel is counterintuitive since Nr6a1 is a well-characterised direct repressor of *Oct4* in ESCs (Gu et al., 2005, 2011; Mullen et al., 2007; Sato et al., 2006) and, consistent with this view, *Oct4* expression is enhanced in the posterior growth zone of Nr6a1 null embryo at E8.5 (Fuhrmann et al., 2001) and in our equivalent-stage *Nr6a1*^-/-^ ESC-derived NMPs (Day2.5 and Day3 NMPs; Figure S10C and D). Why then does *in vivo* deletion of Nr6a1 reduce trunk vertebral number if *Oct4* expression is heightened? We suggest the explanation for this result lies in the timing of gene regulatory mechanisms. While loss of Nr6a1 upregulates *Oct4* within trunk-forming progenitors (E8.5 *in vivo*, Day2.5/3 *in vitro*), it is known that ectopic Oct4 activity has little to no phenotypic consequence within its normal (cervical-thoracic) domain of activity (Aires et al., 2016). It is only at later stages, when endogenous *Oct4* is normally extinguished, that persistent/exogenous Oct4 acts to drive trunk elongation (Aires et al., 2016). Extending our *in vivo* analysis to these later stages showed that *Oct4* transcripts were undetectable in both wildtype and *TCreER*^T2^;*Nr6a1^flx/flx^* tailbuds at E9.5 by qPCR (data not shown). Similarly *in vitro, Oct4* expression dropped to negligible levels in both wildtype and *Nr6a1*^-/-^ cells on d4 of differentiation (Figure S10C), indicating that the upregulation of *Oct4* following loss of Nr6a1 is transient. *Gdf11* and its signalling pathway components became robustly expressed with or without Nr6a1 (*In vitro*: d4 and d4G, Fig S10C; *In vivo*: *TCreER*^T2^;*Nr6a1^flx/flx^*, Fig 2F), and thus would be expected to clear ectopic *Oct4* expression even in Nr6a1 loss-of-function settings, while at the same time as amplifying precocious posterior *Hox* gene expression, collectively reinforcing the premature termination of trunk formation.

In parallel with its role in controlling axial elongation and vertebral number, this work has identified Nr6a1 as a critical intrinsic regulator of *Hox* cluster temporal progression and body patterning. At a mechanistic level, Nr6a1-dependent regulation of *Hox* cluster expression could be indirect via secondary signal(s), particularly as all 4 clusters are concurrently affected. However, reanalysis of chromatin immunoprecipitation studies characterising mouse mesenchymal stem cells *in vitro* (Gurtan et al., 2013) has identified putative Nr6a1 binding sites within each of the four *Hox* clusters, raising the possibility of a more direct mode of regulation. To date, Nr6a1 has been characterised as a transcriptional repressor (Cooney et al., 1998; Yan and Jetten, 2000) which, in the context of the *Oct4* promoter in ESCs, recruits the methyl transferases such as Dnmt3a, Mdb2 and Mdb3 to silence expression (Gu et al., 2011; Sato et al., 2006). Additionally, Nr6a1 has been shown *in vitro* to interact with the histone deacetylase complex component Nuclear receptor corepressor 1 (Ncor1) (Yan and Jetten, 2000) and Ubiquitin interacting motif containing 1 (UIMC1) (Yan et al., 2002) in the repression of downstream target genes. Regarding the latter, the particular SNP identified within Nr6a1 as associated with additional trunk vertebrae in pigs was shown to enhance binding between Nr6a1 and Ncor1/UIMC1 (Mikawa et al., 2007), supporting the potential for a direct mode of Nr6a1 regulation *in vivo*. While mechanism remains to be elucidated, the global effect of Nr6a1 in regulating the temporal activation of expression from all *Hox* clusters supports an ancient role for this nuclear receptor, at least back to the common ancestor of vertebrates when a single cluster existed.

A particularly striking and unexpected aspect of Nr6a1 function revealed in these studies was its axially-restricted role in segmentation. Thoraco-lumbar segmentation required the presence of Nr6a1, though was resistant to exaggerated Nr6a1 levels. Conversely, the active clearance of Nr6a1 function was essential in allowing normal post-sacral segmentation. This switch in requirement for Nr6a1 activity in either scenario was incredibly sharp, visualised both in terms of somite and vertebrae morphology, supporting a very rapid transition in the genetic networks driving this developmental process. Axially-restricted contribution to pre-sacral or post-sacral segmentation has been observed in a small number of studies, highlighting potential downstream effectors of Nr6a1 in this context. For example, some *Hox* genes have been shown to cycle in the PSM (Zákány et al., 2001) and, while altered *Hox* expression or Hox function has not generally not been linked to defects in segmentation, the *in vivo* ectopic expression of a linker region-deficient Hoxb6 protein was shown to cause severe defects in segmentation posterior to the hindlimb only (Casaca et al., 2016). In the case of Nr6a1 deficiency, Hox protein function is not expected to be altered, though it remains possible that aberrant levels of Hox activity may impact segmentation. A second, and more likely, mediator is Lunatic Fringe (Lfng), a core component/inhibitor of the Notch signalling pathway. Throughout segmentation, *Lfng* is expressed both in an oscillatory pattern in the posterior PSM and stably at the anterior PSM border (Evrard et al., 1998; Zhang and Gridley, 1998), controlled by independent enhancers (Cole et al., 2002; Morales et al., 2002). The specific abolition of oscillatory *Lfng* expression via enhancer deletion resulted in highly malformed vertebrae of reduced number specifically within the presacral region, with the tail extending largely unaffected (Shifley et al., 2008; Stauber et al., 2009; Williams et al., 2014). This regional phenotype was very similar to what was observed in *TCre:Nr6a1^-/-^* mutant embryos (Figure 2F), and moreover, cyclic expression of *Lfng* has been found to be greatly reduced in *Nr6a1*^-/-^ embryos while the stable band of *Lfng* in anterior PSM was unaffected (Chung et al., 2001). Collectively, this supports the oscillatory expression of *Lfng*, downstream of Nr6a1, as essential for presacral somite segmentation.

While we have focused on the paraxial mesoderm and its major derivative, it is clear that the entire trunk region expands or contracts following manipulation of Nr6a1 level, suggesting that cell-autonomous Nr6a1 actions are likely to extend across multiple tissue/germ layers in addition to more global mechanisms of tissue-level coordination. For the lateral plate mesoderm, a change in hindlimb positioning that mirrors the anterior or posterior displacement of sacral vertebrae has been observed in Gdf11 signalling gain- and loss-of-function mouse mutants (Jurberg et al., 2013; McPherron et al., 1999) and also here in *Cdx2P:Nr6a1* gain-of-function embryos. In *TCre:Nr6a1^-/-^* embryos however, the positioning of rudimentary pelvic bones maintained an almost “wildtype” distance from the last formed rib-bearing thoracic element, despite all intervening elements being transformed from lumbar to sacral identity. This suggests a likely dissociation in Nr6al’s regulation of *Hox* signatures between tissue layers, and in the downstream events they instruct (Moreau et al., 2019). From an evolutionary perspective, our work provided experimental support for changes in Nr6a1 activity underlying the modest changes in axial formulae observed in domesticated pigs and sheep. Whether analogous changes are widespread across the vertebrates and whether more dramatic changes in expression or function may have supported the evolution of extreme phenotypes such as the elongated vertebral column in snakes remain important areas for future investigation.

## Supplementary Table

**Table S1**

ToppGene and DAVID analysis of *Nr6a1*^-/-^ embryos compared to controls.

## Methods

### Resource availability

Further information and requests for resources and reagents should be directed to and will be fulfilled by Edwina McGlinn.

### Experimental models and subject details

#### Animal models and ethical approval

All animal procedures in Monash University were performed in accordance with the Australian Code of Practice for the Care and Use of Animals for Scientific Purposes (2013). These experiments were approved by the Monash Animal Ethics Committee under project number 21616. Animal experiments performed at the Stowers Institute for Medical Research were conducted in accordance with an Institutional Animal Care and Use Committee approved protocol (IACUC #2019-097).

#### Mice

*Gdf11*^-/-^ (McPherron et al., 1999), *Nr6a1^-/-^* (Chung et al., 2001), *Nr6a1^flx/flx^* (Lan et al., 2003), *CMVCre* (Sladitschek and Neveu, 2015) and *TCreER*^T2^ (Anderson et al., 2013) mouse lines have been previously described, and were maintained on a C56BL/6 background. *TCreER*^T2^ and *Nr6a1^flx/flx^* lines were intercrossed to generate *TCreER*^T2^;*Nr6a1*^+/flx^ males, which were subsequently bred with *Nr6a1^+/f^*^lx^ or *Nr6a1^flx/f^*^lx^ dams for time-mate collection of embryos. A similar breeding strategy was employed to incorporate the *Gdf11*^+/-^ allele onto this *TCreER*^T2^;*Nr6a1*^+/flx^ compound line. Conditional deletion of *Nr6a1* was performed by intraperitoneal administration of Tam (60mg/kg) into the pregnant dam at the specified embryonic stages. *Cdx2P:Nr6a1* transgenic embryos were generated by pronuclear injection by the Monash Gene Modification Platform. The full coding sequence of *Nr6a1* long isoform was synthesised with Notl restriction enzyme site addition at the 5’ and 3’ ends by Biomatik, USA, sequence verified, digested, and cloned downstream of the Cdx2P promotor (Benahmed et al., 2008). Two adult transgene-positive chimeras were recovered from multiple rounds of injection, however, a stable line could not be produced as one founder was infertile and one did not transmit the transgene. Thus transient transgenic embryos were collected at E10.5 for tailbud RNA extraction and/or *in situ* hybridisation, and at E18.5 for skeleton preparation. Transgene copy number for each embryo was confirmed by droplet digital PCR.

#### Genotyping

For routine genotyping, tail tissue was collected from postnatal pups and yolk sac collected from embryonic samples. Genotyping was performed by Transnetyx or in-house. For detection of *Nr6a1* conditional deletion via *TCreER*^T2^ and Tam administration at E7.5 or E8.5, digits of the left hindlimb were collected based on known pattern of expression of the T promoter (Anderson et al., 2013; Perantoni et al., 2005). To isolate the DNA, digit tissue was treated with Proteinase K (20µg/ml) in 500µL of Tris-HCl buffer (Composition: 100mM Tris-HCl (pH8.0); 5mM EDTA; 200mM NaCl; 0.2% SDS) overnight at 56C. The DNA was precipitated by adding equal volume of isopropanol and pelleted at 13,000 rpm. Next, the pellet was washed twice with 70% ethanol and eluted in nuclease-free water. At this point, genotyping was performed by standard PCR method using primers described in (Lan et al., 2003).

#### ESCs maintenance

A mouse Bruce-4 ESC line was used for all *in vitro* differentiation assays. ESCs were routinely maintained on an in-house generated mitotically inactive primary mouse embryo fibroblast feeder layer in ES medium (81.8% Knockout DMEM (Gibco, 10829-018); 15% gamma irradiated fetal bovine serum (Gibco, 10101-145); 1% Pen/Strep (Gibco, 15140-122); 1% GlutaMAX-I (Gibco, 35050-061); 1% MEM NEAA (Gibco, 11140-040), 0.2% 2-Mercaptoethanol (Gibco, 21985-023)) supplemented with LIF (1000x, made in-house).

#### Generation of *Nr6a1*^-/-^ ESC line

*Nr6a1* null allele generation was performed by the self-cloning CRISPR/Cas9 (scCRISPR) method (Arbab and Sherwood, 2016; Arbab et al., 2015). Two protospacers in *Nr6a1* exon4 were determined using the tool, CRISPOR (http://crispor.tefor.net/) (Concordet and Haeussler, 2018). The sgRNA oligonucleotides order for scCRISPR contains the protospacer (19-21bp), flanked by sgPal7 (Addgene, 71484) homology (U6 promoter and sgRNA) sequence, total 60bp. Three consecutive PCR steps were performed using three pairs of standard primer sets (sgRNA_HDRstep1-3) for preparing homology directed repair (HDR). ESCs used for electroporation were cultured and maintained as previously described without antibiotics. Once the ESCs were ready, they were pelleted and suspended in buffer R (ThermoFisher, MPK1025) at a density of 1*10^7^ cells/ml. 100µl of the ESCs were then mixed with the scCRISPR construct (Composition: 10µl elute of HDR PCR product; 1µl sgPal7 (Addgene, 71484); 1µl spCas9-BlastR (Addgene, 71489). ESCs were electroporated at pulse voltage 1,400 and width (ms) 3 for three cycles, and then plated on a 10 mm feeder-covered plate. 96 single colonies were selected and expanded independently in a 96-well plate. Multiple independent knockout lines were produced by this method and clones selected based on normal morphology. Clone B1 used in these studies harboured independent genomic deletions within exon 4 of each *Nr6a1* allele, one deleting l25bp (Δl25) and the second 2l9bp (Δ2l9) (Figure S8). Compared to the wildtype Nr6a1-long isoform protein product of 495 amino acids (aa), Δl25 generates a truncated protein of 126 aa with no DNA binding and ligand binding domain, while the in-frame deletion Δ2l9 allele encoded a protein of 422 aa with no DNA binding domain, both supportive of a functional null (Lan et al., 2002).

### Method details

#### Gene cloning

*Nr6a1* isoform sequences were confirmed using gene amplification from E10.5 embryonic tissue. RNA was extracted (Macherey-Nagel) and cDNA prepared (Roche, 04897030001) using both random hexamers and oligodT primers. Sequence-specific primers were designed based on the version mm10 genome (UCSC) as detailed in Key resources table. Amplified sequences were cloned into pGEM-T easy vector (Promega) and used as the template for synthesising RNA riboprobes.

#### Whole mount *in situ* hybridisation

Whole-mount *in situ* hybridisation was performed as previously described (McGlinn et al., 2019) with minor modifications. Mouse embryos were dissected and placed in iced-cold PBS (Gibco, 14190-144). For embryos E9.5 or older, the brain’s 4^th^ ventricle was pierced to prevent probe trapping. Embryos were fixed in 4% paraformaldehyde (PFA) at 4C, rocking overnight. Fixed embryos were washed twice in PBT (Composition: 137mM NaCl; 2.7mM KCl; 10mM Na_2_HPO_4_; 2mM KH_2_PO_4_ in DEPC-H_2_O (pH7.4); 0.1% Tween) for 5 min and dehydrated into methanol using a graded MeOH/PBT series (25%, 50%, 75%, 100% MeOH) for 5-20 mins each with gentle rocking at room temperature (RT). To start *in situ* hybridisation, embryos were rehydrated into PBT using a graded MeOH/PBT series (75%, 50%, 25% MeOH) for 5-20 mins each and washed in PBT twice for 5 min with gentle rocking at RT. The embryos were then treated with 10 µg/ml of proteinase K in PBT as follows: E8.5 for 3.5 min, E9.5 for 8 min, E10.5 for 15 min and E12.5 for 25 min at RT with gentle rocking. The reaction was quenched by washing embryos twice in PBT for 5 min and following postfix in 4% PFA with 0.2% glutaraldehyde for 20 min at RT. Embryos were then washed twice in PBT for 5 min at RT and transferred to hybridisation solution (Composition: 50% formamide; 5x SSC (pH4.5), 1% SDS; 50µg/ml heparin; 50µg/ml yeast tRNA (Sigma, R6750)) at 70C with constant rocking for at least 2 hr before 1µg/ml DIG-labeled riboprobe was added and incubated overnight. On the second day, the embryos were washed 3 times at 70C in pre-warmed solution I (Composition: 50% formamide; 5x SSC (pH4.5); 1% SDS) for 30 min, followed by 3 washes at 65C in pre-warmed solution II (Composition: 50% formamide; 2x SSC (pH4.5); 0.1% Tween-20) with gentle rocking. The embryos were then washed 3 times in TBST (Composition: 137mM NaCl; 2.7mM KCl; 25mM Tris-HCl (pH7.5); 0.1% Tween-20) for 5 min at RT, and blocked in TBST containing 10% heat-inactivated sheep serum (HISS) for 2 hrs at RT with gentle rocking. The embryos were then transferred to TBST containing 10% HISS and anti-DIG-AP antibody (Roche, 11093274910) at a dilution of 1:2000 overnight at 4C with constant rocking. On the third day, the embryos were washed at least 5 times in TBST for l hr each wash at RT with gentle rocking and then overnight in TBST at 4C. On the fourth day, embryos were equilibrated into NTT (Composition: 100mM NaCl, 100mM Tris-HCl (pH9.5); 0.1% Tween-20) by washing 3 times for 10 min each at RT with gentle rocking. To develop the color, the embryos were incubated in BM purple (Roche, 11442074001) at RT. To stop the color reaction, embryos were washed 3 times in PBT for 5 min, and postfixed in 4% PFA for 20 min at RT. The embryos were then washed 3 times in PBT for 5 min and stored in PBT at 4C until photographing.

#### Microarray

E9.0-E9.5 wildtype and *Nr6a1^-/--^* embryos were isolated and yolk sac DNA genotyped in parallel. Total RNA was extracted from each whole embryo using TRIzol reagent and processed through an RNeasy column (QIAGEN) using the RNA clean-up protocol. Concentration and quality of RNA were determined by spectrophotometer and Agilent bioanalyzer analysis (Agilent Technologies, Inc., Palo Alto, CA). For array analysis, labeled mRNA targets were prepared from 150 ng total RNA using MessageAmp III RNA Amplification Kit (Applied Biosystems / Ambion, Austin, TX) according to manufacturer specifications. Array analysis was performed using Affymetrix GeneChip GeneChip Mouse Genome 430 2.0 Arrays processed with the GeneChip Fluidics Station 450 and scanned with a GeneChip Scanner 3000 7G using standard protocols.

#### Tailbud dissection for gene expression analysis

For E9.5 tailbud collection, all tissue at and caudal to the posterior neuropore was collected. For E10.5 tailbud collection, all tissue caudal to the last somite was collected. Dissected tissue was immediately placed into lysis buffer (Macherey-Nagel) on dry ice and stored at -80C. RNA was extracted (Nucleospin RNA kit, Macherey-Nagel, 740955). The remaining part of embryos were processed for whole-mount *in situ* hybridisation detection of *Uncx4.1* and accurate somite count determination.

#### Quantitative PCR using BioMark Fluidigm

100 ng RNA isolated from E9.5 and E10.5 tailbud tissue was used as template for cDNA synthesis performed using RT-Vilo (ThermoFisher). Quantitative PCR was performed using the 96*96 BioMark Fluidigm format. Raw Ct values were analysed using a modified version of the qPCR-Biomark script (https://github.com/jpouch/qPCR-Biomark) and normalised as previously described (Livak and Schmittgen, 2001). Only Ct-values in the optimal range for the Biomark system of 6-25 were used for further analysis. All genes were first normalised against the mean raw Ct-values of five housekeeping gene probes yielding ΔCt values, then normalised against the wildtype condition yielding ΔΔCt values.

#### Quantitative PCR using Roche Lightcycler

RNA isolated from one E9.5 tailbud or 1mg RNA isolated from the *in vitro* cells was used as template for cDNA synthesis (Roche, 04897030001) using random hexamers and eluted in 200µl and 100µl H_2_O respectively. Each 10µl PCR reaction reaction (5µl SYBR Green I Master Mix (Roche, 04887352001); 2.5µl H_2_O; 2µl cDNA; 0.25µl 10mM forward primer; 0.25µl l0mM reverse primer) was run on a Lightcycler 480 (Roche) using the program: 95C for l0s (l cycle), 95C for l0s, 60C for l5s, 72C for l0s (45 cycles). Biological samples were processed in technical triplicate for each gene and the expression of housekeeping gene *Pol2a* used for normalising gene expression using the ΔΔCt method.

#### RNA-seq

For *in vivo* sample RNAseq, tissue was dissected and RNA extracted as described above. RNA quality and quantity was assessed using the Agilent Technologies BioAnalyser. Multiplexed RNAseq libraries, generated from 50ng total RNA per sample, were prepared using Illumina TruSeq Stranded mRNA Sample prep (Protocol 15031047 Rev Oct 2013). 75bp single-end (SE) sequencing was performed on the NextSeq500 Illumina platform. On average, 69 million SE reads were obtained for each library. For *in vitro* cell sample RNAseq, libraries were prepared by an in house method. The index is added during initial pA priming and pooled samples amplified using template switching oligos. P5 was added by PCR and P7 by Nextera transposase. The final library structure is as follows: P5-Rd1->8bp index-10bpUMI-pA then cDNA<-Rd2 primer i7 index P7. Paired end sequencing was performed on the NextSeq550 Illumina platform for 19bp forward reads of index and 72bp reverse reads of cDNA. On average, 20M raw reads were obtained for each library.

#### Skeleton preparation and imaging

Skeletal preparation was performed on E16.5 and E18.5 embryos as previously described (McLeod, 1980). Embryos were skinned, then incubated for 2 days at RT with constant rocking in each of the following solutions: 95% ethanol, 100% acetone, staining solution (Composition: 15 mg alcian blue and 5mg alizarin red S in 70% ethanol with 0.5% glacial acetic acid). To clear the skeletons, stained embryos were washed in 1% KOH at RT with constant rocking for 2-5 days, then transferred into glycerol using a graded glycerol/1% KOH series (25%, 50%, 75%, 100% glycerol) for 24hr in each solution with gentle rocking at RT. Finally, the embryos were transferred in 100% glycerol with 0.02% sodium azide for long-term storage. Images were acquired with a Vision Dynamic BK Lab System at the Monash University Paleontology Lab. Images were taken with a Canon 5d MkII with a 100mm Macro lens (focus stop 1:3/1:1). Multiple images were taken to extend the focal depth, and stacked in ZereneStacker using the PMax algorithm.

#### NMP differentiation of ESCs

NMP differentiation was based on the published protocols (Gouti et al., 2014, 2017) with minor modifications. In preparation for differentiation, ESCs were feeder-depleted and plated in gelatin-coated (0.1% Sigma, G1890-100G) 6-well plates (Falcon, 353046) at a density of 8×10^3^ cells/cm^2^ in ES medium. On D0 of differentiation media was changed to N2B27 media (Composition: 49.5% Advanced Dulbecco’s Modified Medium F-12 (Gibco, 12634028); 49% Neurobasal medium (Gibco, 21103049); 0.5% N2-supplement (Gibco, 17502001); 1% B27- supplement (Gibco, 17504044)) supplied with 1x Glutamax (Gibco, 17504044), 40 µg/ml BSA Fraction V (Gibco, 15260037) and 100 mM 2-Mercaptoethanol (Gibco, 21985-023)) supplemented with 10 ng/ml hFGF-2 (Miltenyi Biotec, 130-104-925). The media was further supplemented with 5µM CHIR99021 (StemMACS, 130-103-926) on D2 and with or without 50 ng/ml hGdf11 (130-105-776, Miltenyi Biotec) on D3. Throughout differentiation, the medium was refreshed every 24 hours. NMP identity at D3 was routinely confirmed by co-expression analysis of *Sox2* and *T/Bra* using immunofluorescence.

### Quantification and statistical analysis

#### Microarray analysis

Microarray CEL files were analyzed using RMA (Irizarry et al., 2003) and limma (Smyth, 2004) in the R statistical package. Following alignment and annotation of genes, the Microarray data was filtered to export differentially expressed genes with an adjusted p-value or false discovery rate of 0.05 or lower and a fold change less than or equal to -2 and greater than or equal to 2 (log_2_FoldChange less than or equal to -1 and greater than or equal to 1). Genes were then sorted to extract the top 50 most downregulated genes, which were analysed via DAVID (Huang et al., 2009a, 2009b) and ToppGene (Chen et al., 2009) utilizing standard settings provided in each program. The results were then exported into tables in Excel and the bar graph of biological processes affected by the top 50 downregulated genes was generated using - log10(p-value) parameters and plotted using PRISM.

#### Vertebral formulae

For statistical analysis and data visualisation of vertebral numbers, R-package “Data Analysis using Bootstrap-Coupled ESTimation” (dabestr) was used, and the background of the methods were previously described (Ho et al., 2019). To determine mean differences to the respective shared control, 5000 bootstrap samples were taken and the confidence interval was bias-corrected and accelerated. In the visualisation, 95% confidence interval is indicated by the ends of the vertical error bar and the sampling error distribution is diagrammed as a grey filled curve. The codes are available at https://github.com/ACCLAB/dabestr.

#### Quantitative PCR using Roche Lightcycler

All genes were first normalised against the mean raw Ct-values of housekeeping gene, *Pol2a*, yielding ΔCt values. Following normalisation of gene expression using the ΔΔCt method, statistical analysis was performed using the Wilcoxon test.

#### RNAseq analysis

All sequencing reads were aligned to the reference genome using STAR aligner (Dobin et al., 2013). Only the genes with counts, which are greater than 10, and with CPM, which are greater than 2 in two biological replicates were used for further analysis. Differential gene expression between control and mutants were performed using edgeR (McCarthy et al., 2012; Robinson et al., 2010). Here we used false discovery rate (FDR), the adjusted p value, to display significance of the differential gene expression between the controls and the mutants. Genes with a FDR < 0.05 were considered to be significantly differential expressed in the mutants.

### Key resources table

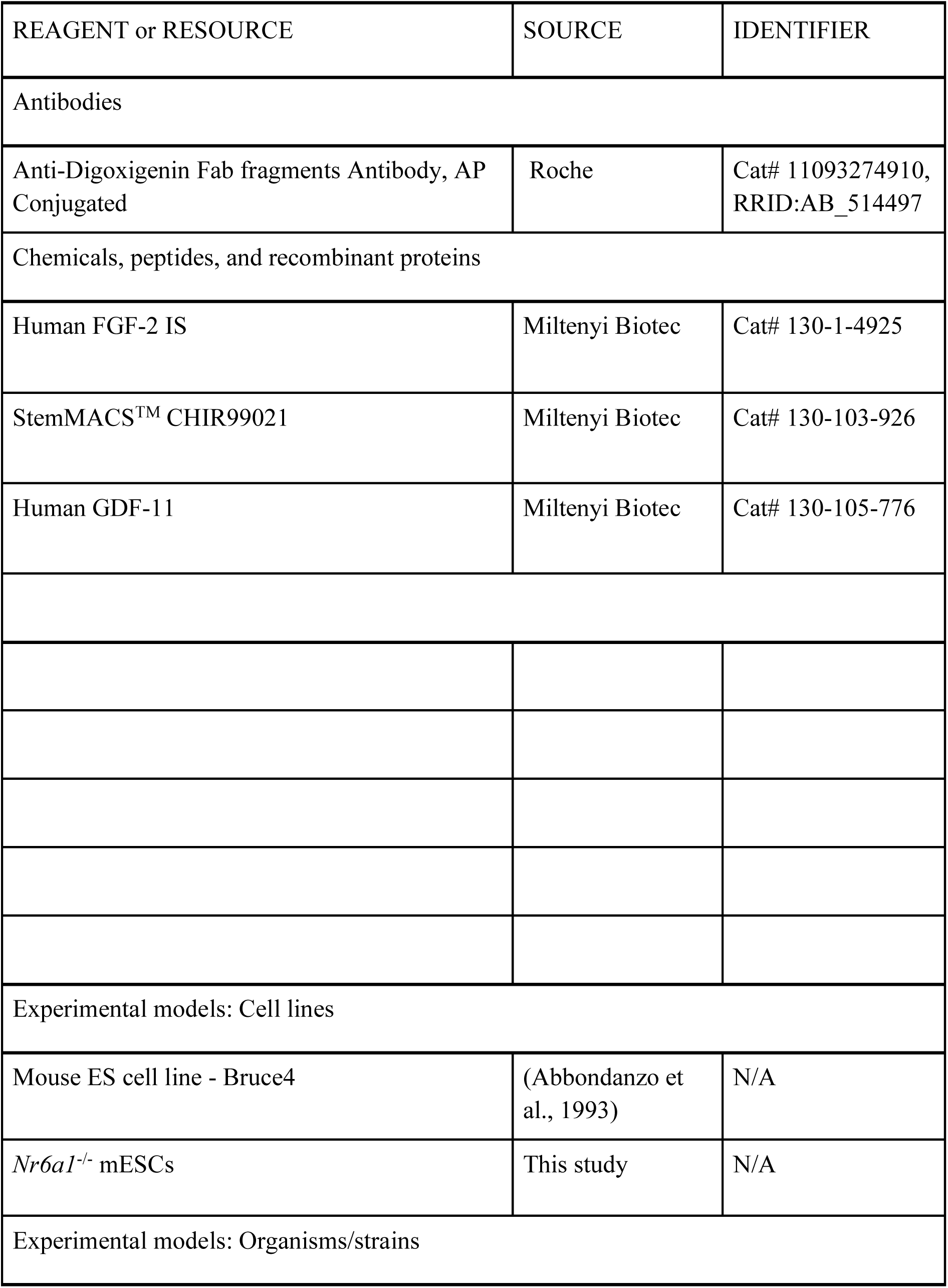
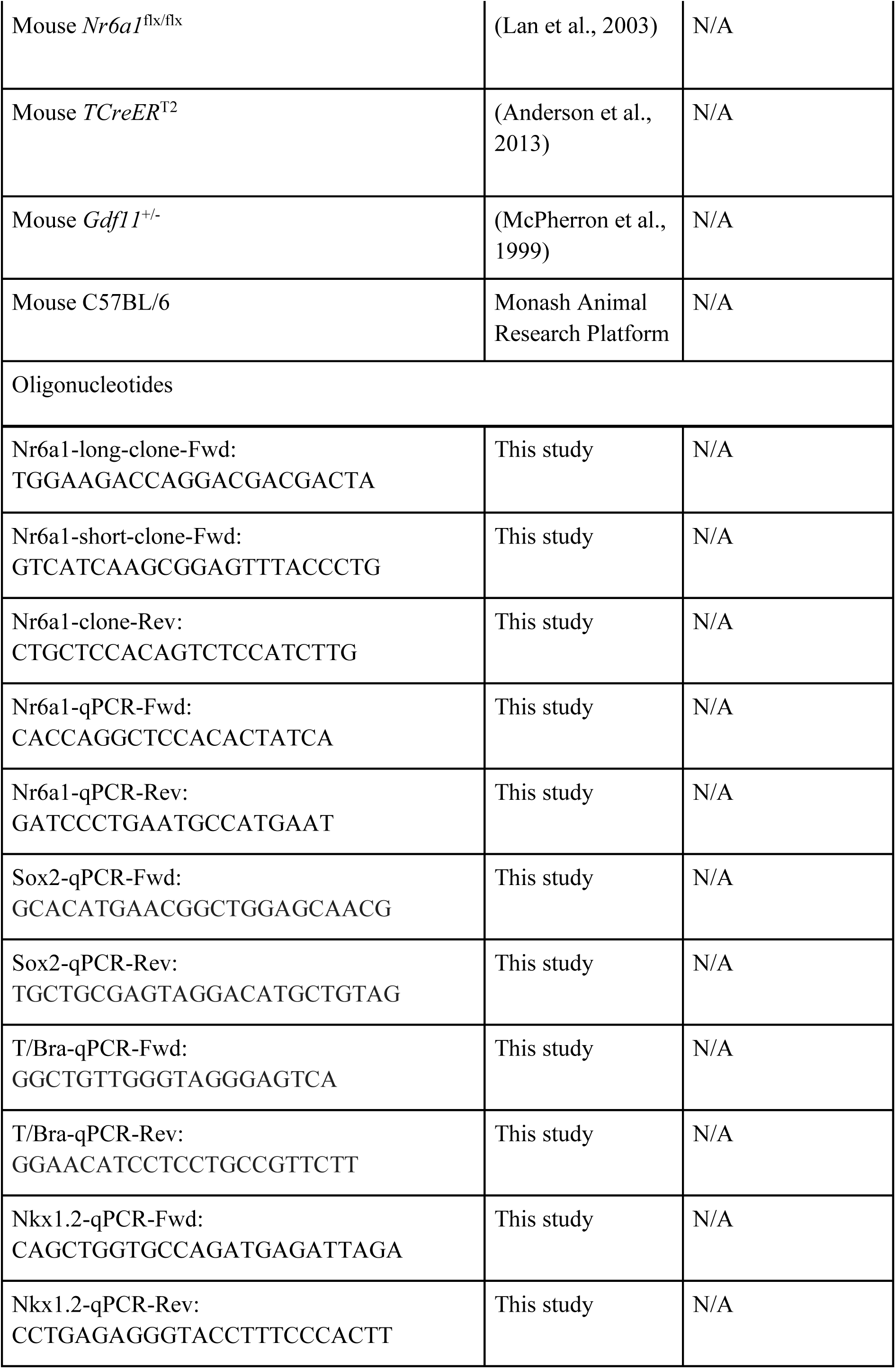
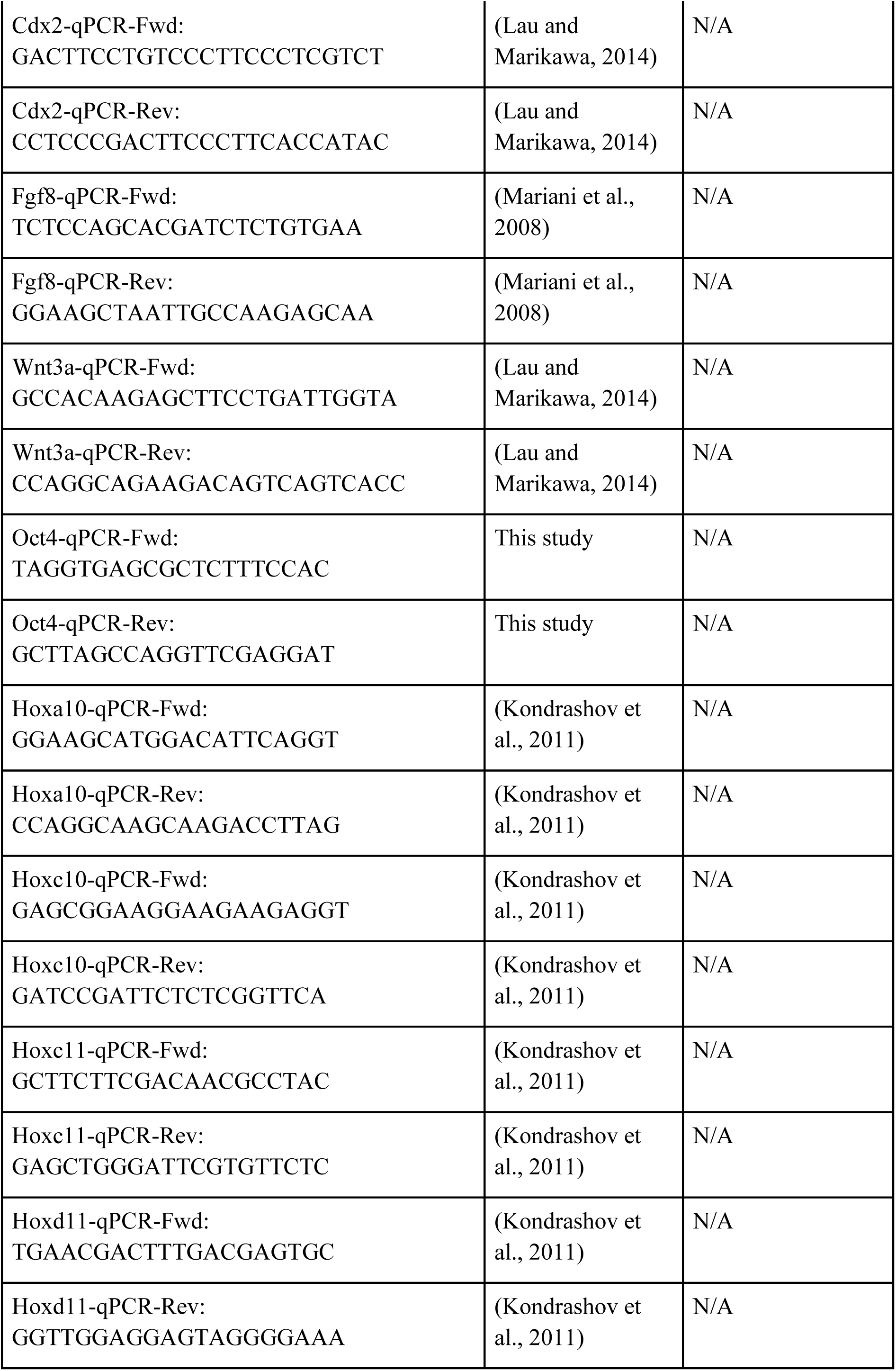
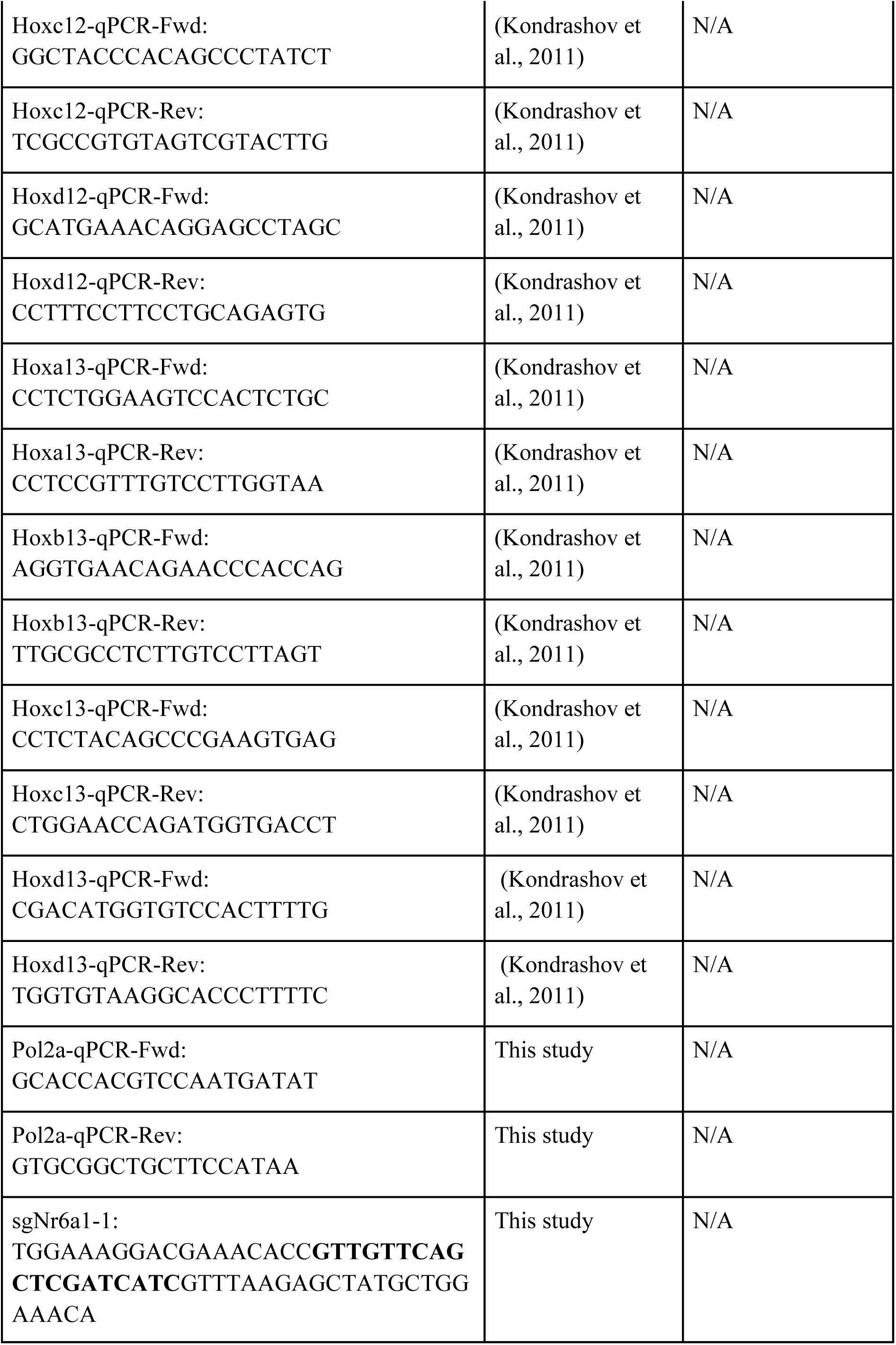
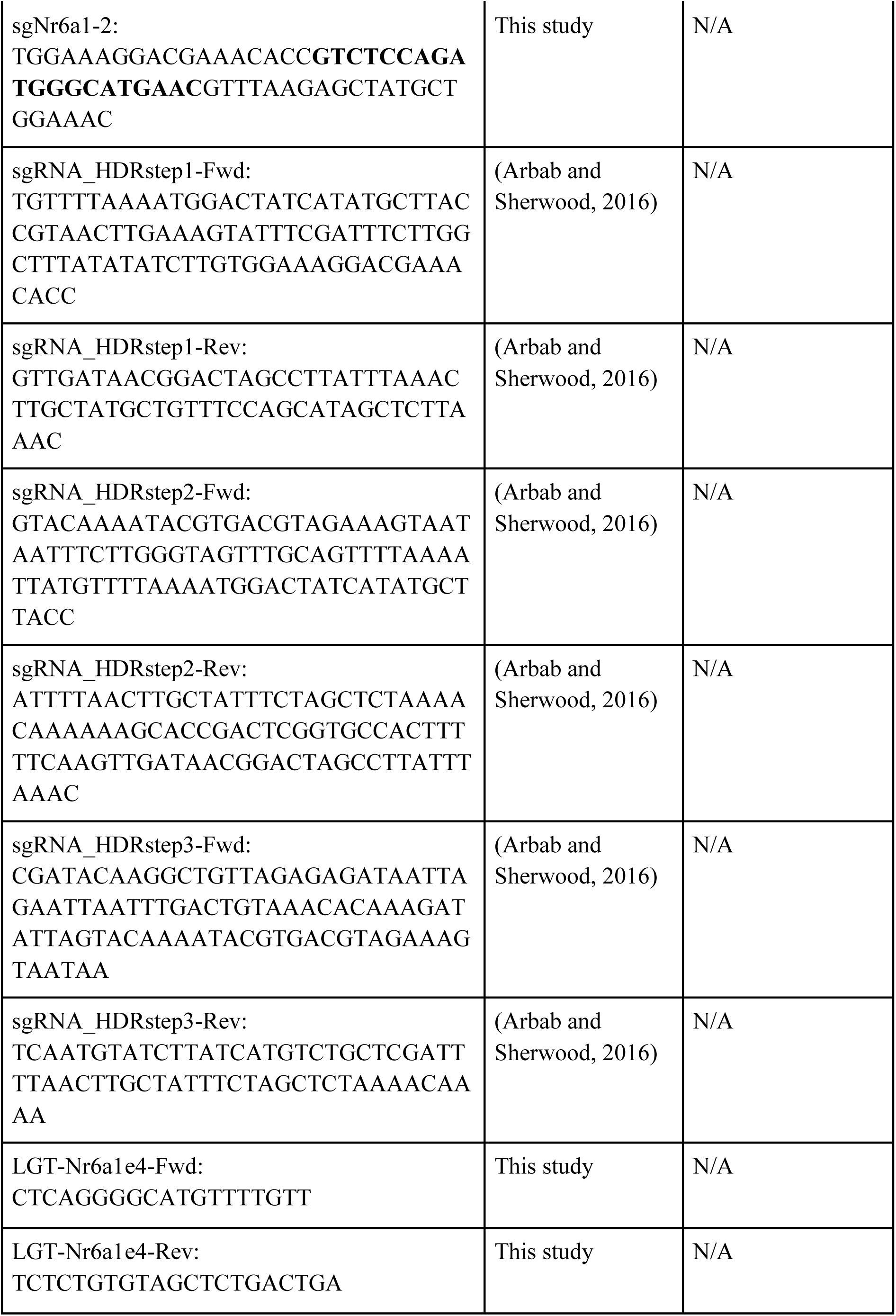
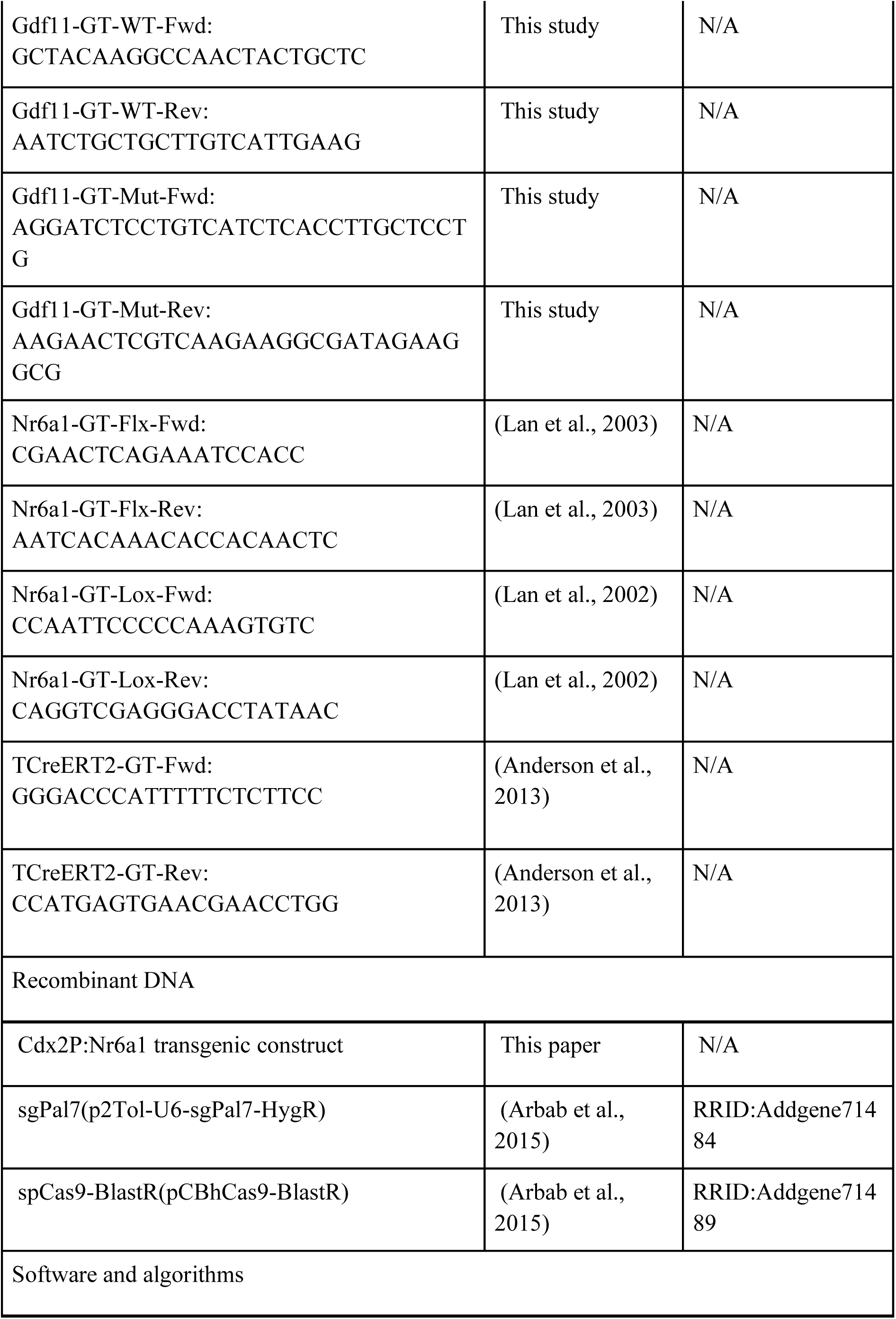
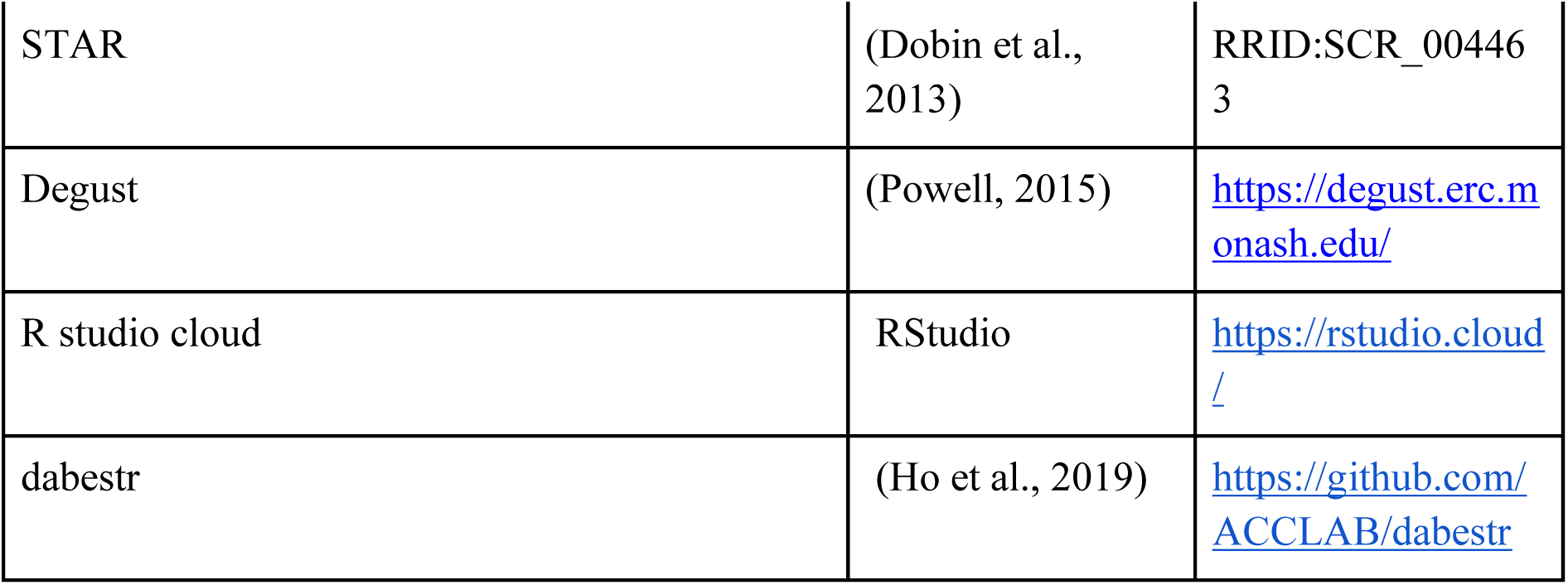

## Acknowledgements

The authors acknowledge the excellent technical assistance provided by Monash Gene Modification Platform, Monash Animal Research Platform and the MHTP Medical Genomics Facility, Karin Zueckert-Gaudenz and Allison Peak for their microarray technical expertise and to Dr Chris Seidel for bioinformatic analysis support. We also greatly appreciate Melissa Childers and the Laboratory Animal Services Facility at the Stowers Institute for their care and maintenance of our mice. The *Gdf11* mutant mouse line was kindly provided by Professor Se-Jin Lee. Y-C.C. and G.M.H. are supported by an Australian Government Research Training Program Scholarship. A.A was the recipient of an American Association for Anatomy Post-Doctoral Fellowship. J.M.P is supported by an Australian Research Council Future Fellowship. This work was supported by Australian Research Council Discovery Project DP180102157 to E.M. and J.M.P. Research in the Trainor laboratory is supported by the Stowers Institute for Medical Research. The Australian Regenerative Medicine Institute is supported by grants from the State Government of Victoria and the Australian Government.

---

**Figure S1.**
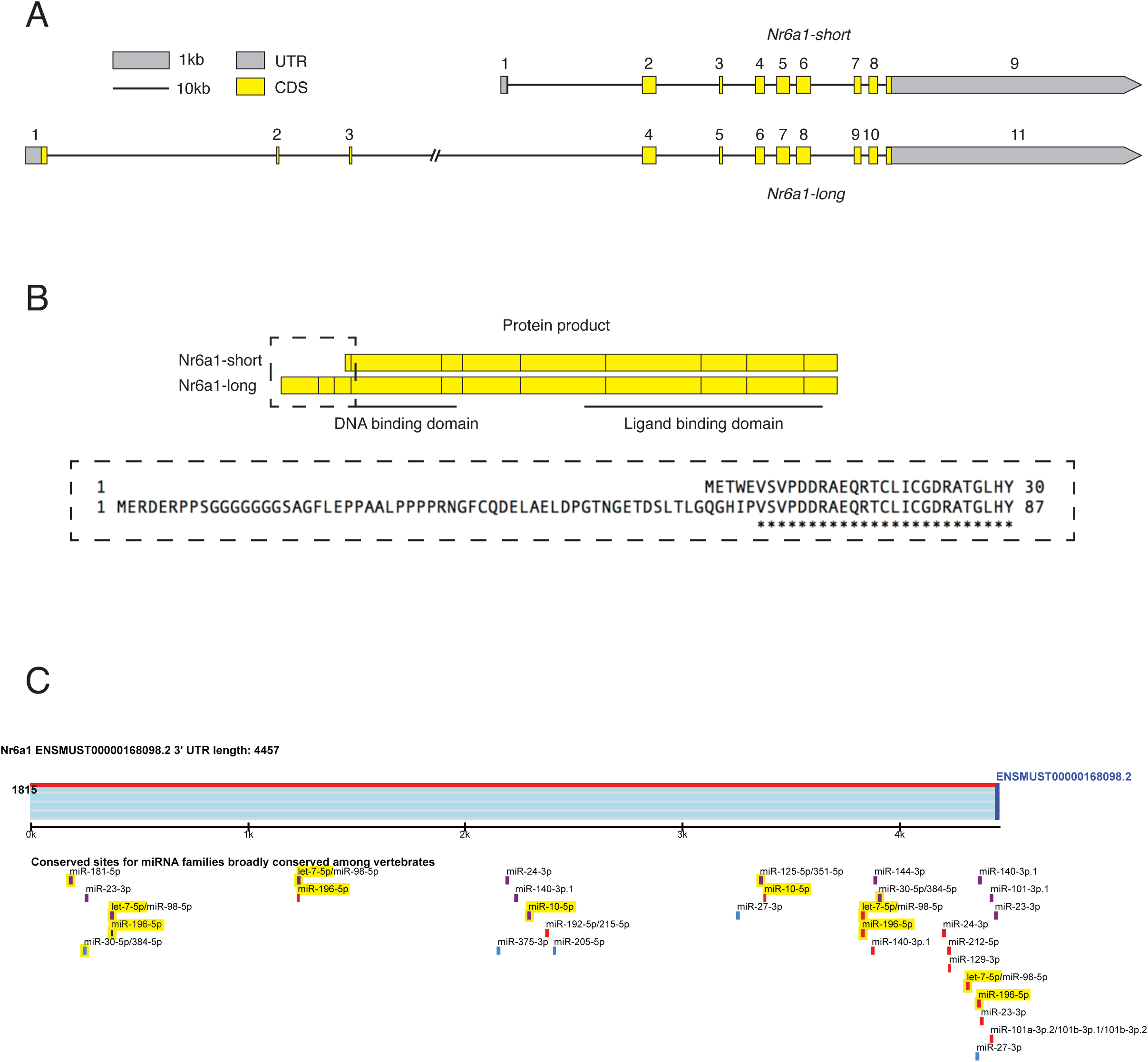
The Nr6a1 genomic locus generates multiple isoforms in the mouse. (A) The genomic structure of *M.musculus Nr6a1* short and long isoforms, drawn approximately to scale, indicating the position of exons (boxes) and introns (lines). Within exons, yellow represents coding sequence (CDS) and grey represents untranslated region (UTR). (B) The protein products (hatched yellow boxes) encoded by *Nr6a1* short and long isoforms share the majority of the DNA binding domain and the entirety of the ligand binding domain. Alternative start sites for each isoform produce variation in the protein sequence of the N-terminus as shown. (C) MicroRNA binding sites located within the *Mus musculus Nr6a1* 3’ UTR, identified using TargetScan (Agarwal et al., 2015). The binding sites of two *Hox*-embedded microRNAs, *miR-10* and *miR-196* were highlighted in yellow.

**Figure S2.**
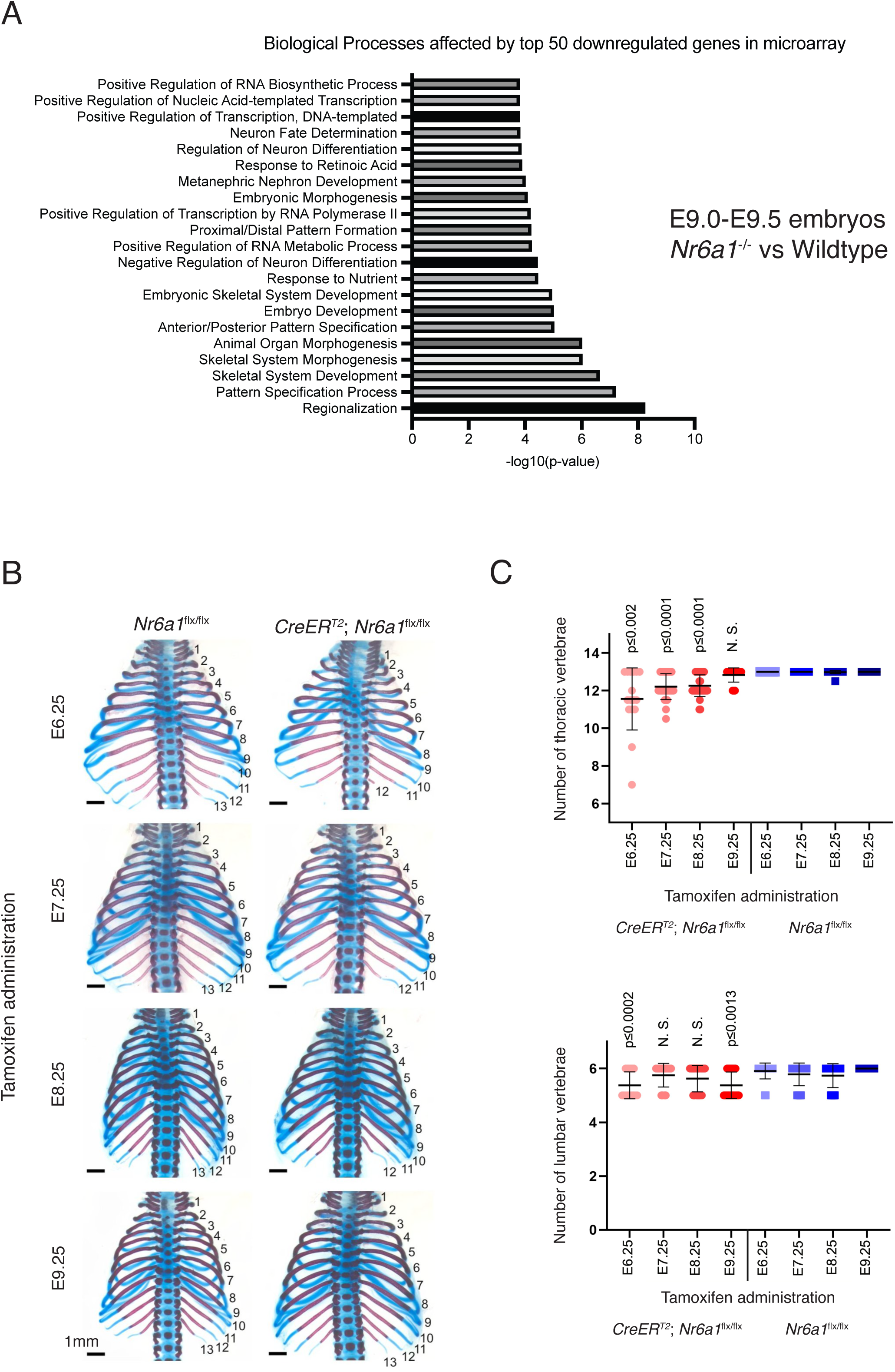
Nr6a1 is required prior to E9.25 to control thoracic vertebral number. (A) The predicted key biological processes disrupted in Nr6a1^-/-^ mutants compared to wildtype controls, which was analysed by ToppGene using the top 50 down-regulated genes in Nr6a1^-/-^ mutants (Chen et al., 2009). (B) Characterisation of the ubiquitous but temporally-controlled deletion of Nr6a1, achieved by crossing *CreER^T2^* and *Nr6a1^flx/flx^* lines and Tamoxifen administration to pregnant dams on a single day of development (E6.25-E9.25) as indicated. E=embryonic day. Skeletal preparation of E18.5 *CreER^T2^*;*Nr6a1^flx/flx^* embryos revealed a reduction in the number of rib-bearing thoracic elements compared to *Nr6a1^flx/flx^* when Tamoxifen was administered prior to E9.25. Scale bars are 1mm. (C) Quantification of the number of thoracic and lumbar elements in *CreER^T2^*;*Nr6a1^flx/fl^* and *Nr6a1^flx/fl^* embryos across all timepoints analysed. Un-paired t-test was used for comparisons between mutant and control embryos under the same treatment.

**Figure S3.**
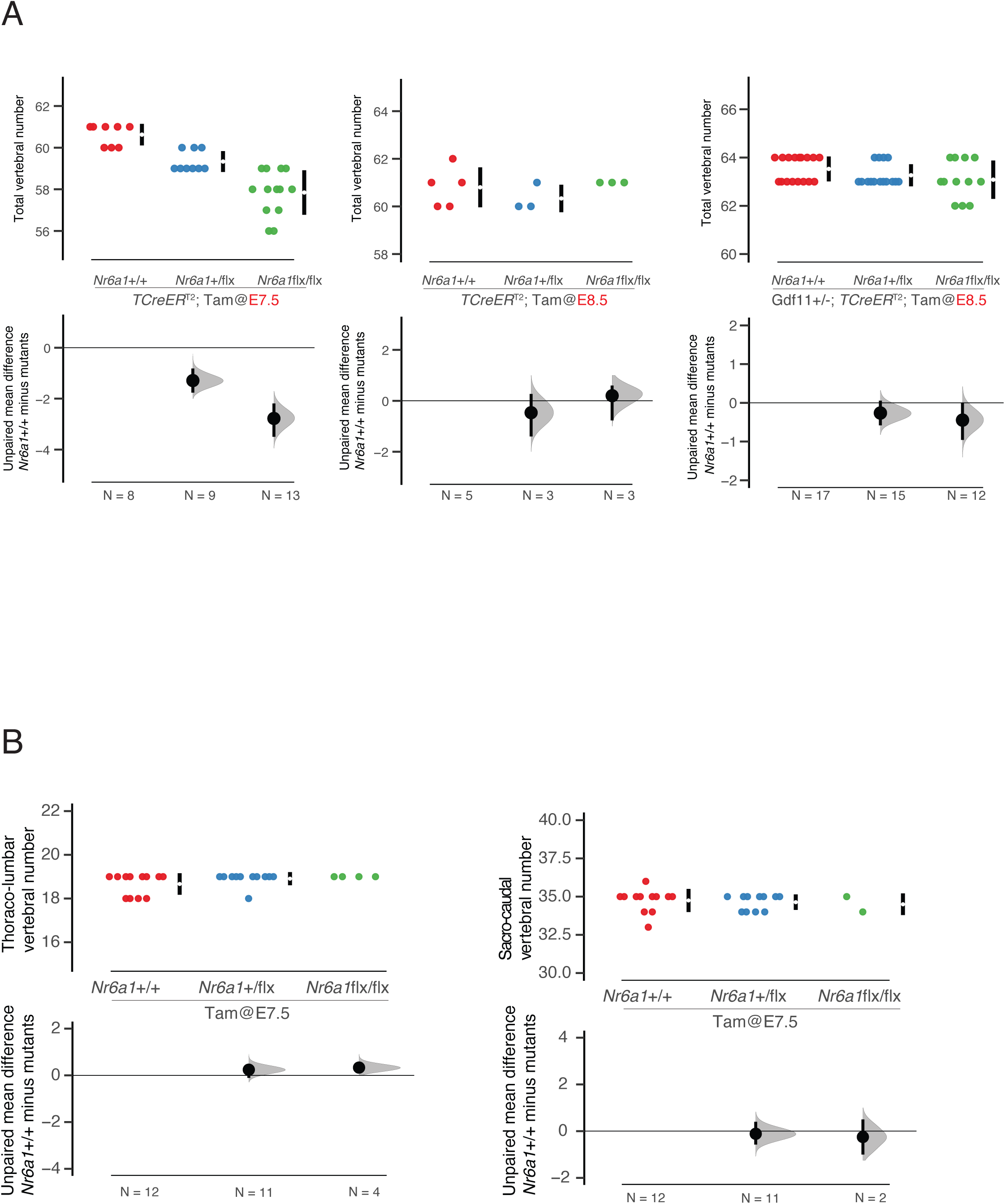
Quantification of vertebral alterations following TCreER^T2^-mediated deletion of Nr6a1. (A) Quantification of total vertebral number following *TCreER*^T2^-mediated conditional deletion of *Nr6a1*. Tamoxifen (Tam) administration to pregnant dams at E7.5 resulted in a dose-dependent reduction in total vertebral number compared to wildtype, while Tam administration at E8.5 resulted in no difference between genotypes. Raw data is presented in the upper plots. Mean differences relative to *Nr6a1*^+/+^ are presented in the lower plots as bootstrap sampling distributions. The mean difference for each genotype is depicted as a black dot and 95% confidence interval is indicated by the ends of the vertical error bar. (B) As a control, Tam administration at E7.5 in TCreER^T2^-negative animals showed no difference in thoraco-lumbar or sacro-caudal count. Raw data is presented in the upper plots. Mean differences relative to *Nr6a1*^+/+^ are presented in the lower plots as bootstrap sampling distributions. The mean difference for each genotype is depicted as a black dot and 95% confidence interval is indicated by the ends of the vertical error bar.

**Figure S4.**
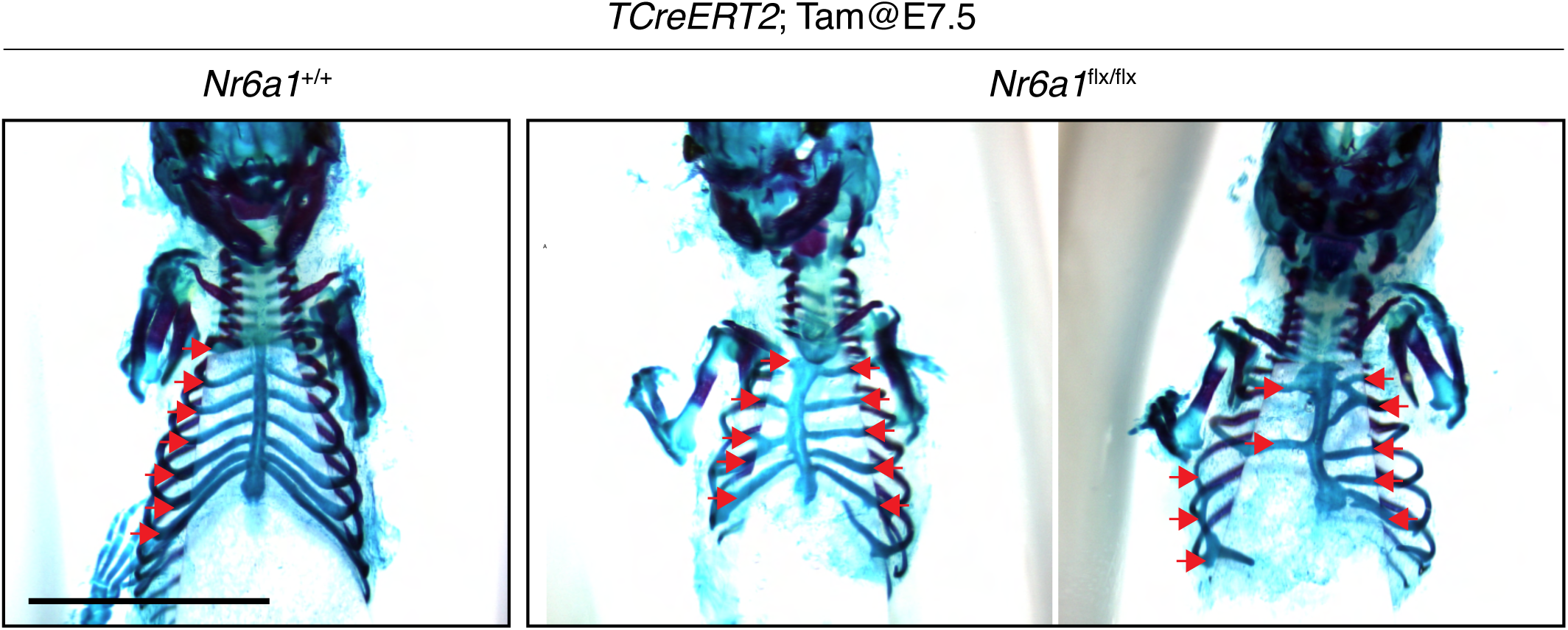
Sternal rib reduction and malformation observed in *TCreER*^T2^;*Nr6a1*^flx/flx^ embryos. Skeletal preparation of E16.5 embryos, ventral view, revealed a reduction in the number of sternal ribs (red arrows) and widespread rib fusions and malformations in *TCreER*^T2^;*Nr6a1^flx/flx^* embryos compared to wildtype. Tamoxifen (Tam) administration at E7.5.

**Figure S5.**
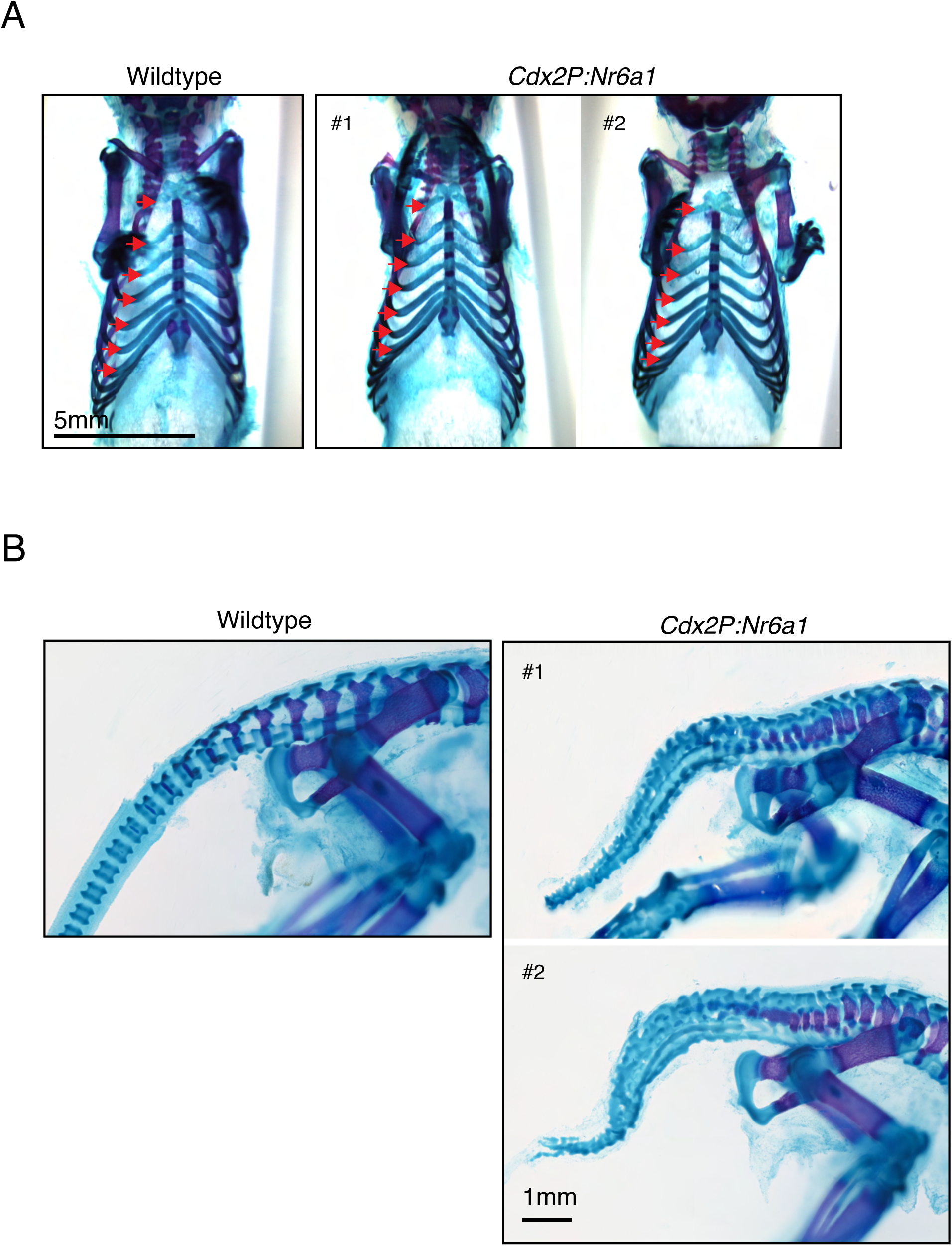
Sternal rib and tail phenotype observed in *Cdx2P:Nr6a1* transgenic embryos. (A) Skeletal preparation of E18.5 embryos, ventral view, revealed no change in the number of sternal ribs (red arrow) in *Cdx2P:Nr6a1* embryos compared to wildtype. (B) Skeletal preparation of E18.5 embryos, lateral view, revealed a highly dysmorphic tail in *Cdx2P:Nr6a1* embryos, appearing to harbour many small fused elements.

**Figure S6.**
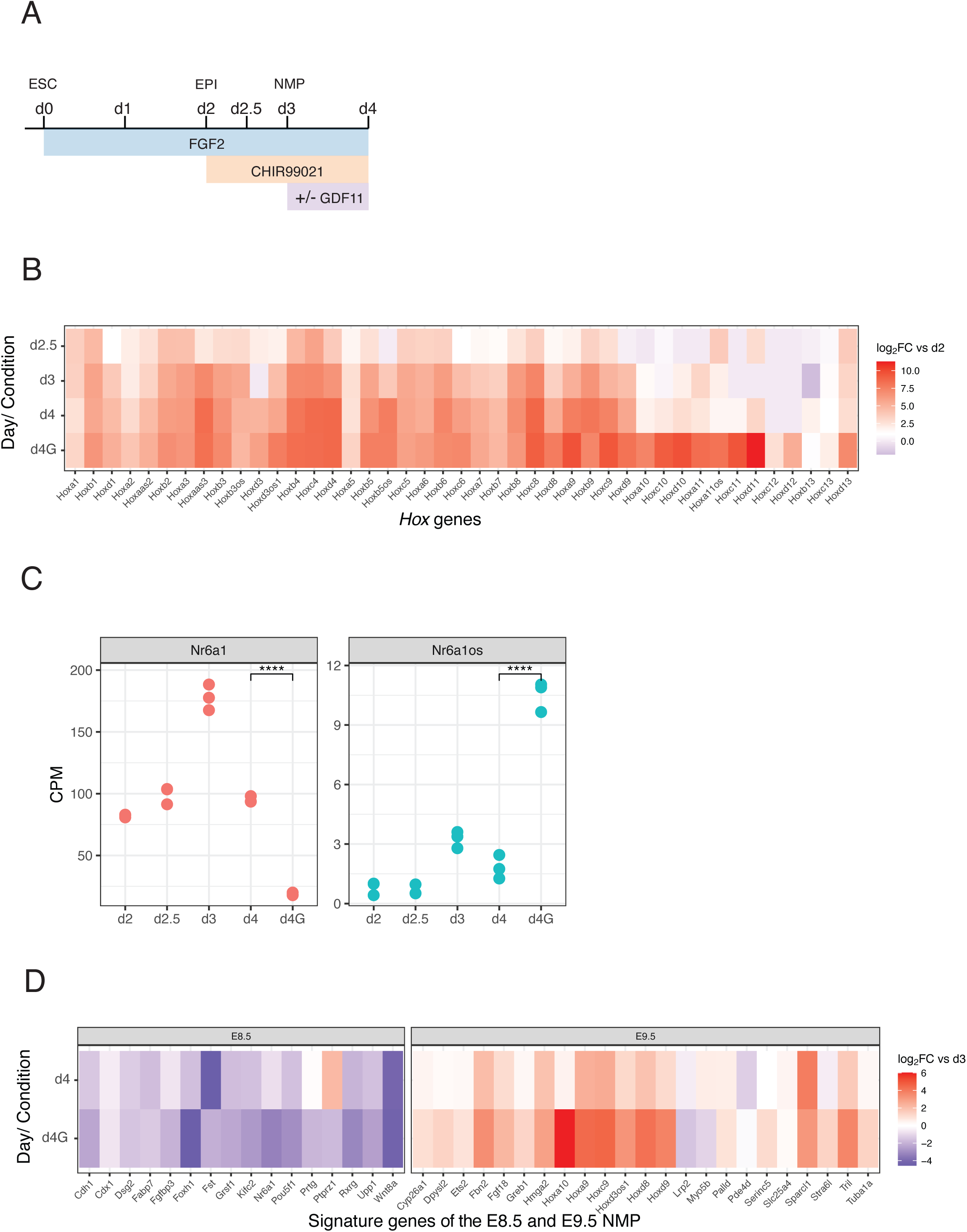
Characterisation of wildtype *in vitro* differentiation kinetics. (A) Schematic representation of the embryonic stem cell (ESC) *in vitro* differentiation protocol for generating epiblast-like (EPI) and neuromesodermal-like (NMP) cells over time. (B) Complete collinear activation of *Hox* genes requires the sequential addition of CHIR99021 and Gdf11. Results are presented as a log2-transformed fold change of expression in d2.5, d3, d4 and d4G cells relative to d2. n= 3 technical replicates per day and per condition. (C) *Nr6a1* and *Nr6a1os* expression analysis over the course of *in vitro* differentiation as schematised. RNA-seq data is presented as counts per million (CPM) and statistical changes in gene expression downstream of Gdf11 addition was performed using edgeR. ****FDR< 0.0001. (D) Temporal change in expression of the temporal signature genes for E8.5 and E9.5 NMPs (Gouti et al., 2017). Results are presented as a log2-transformed fold change in d4 and d4G cells relative to d3 NMPs, n= 3 sample per cell type.

**Figure S7.**
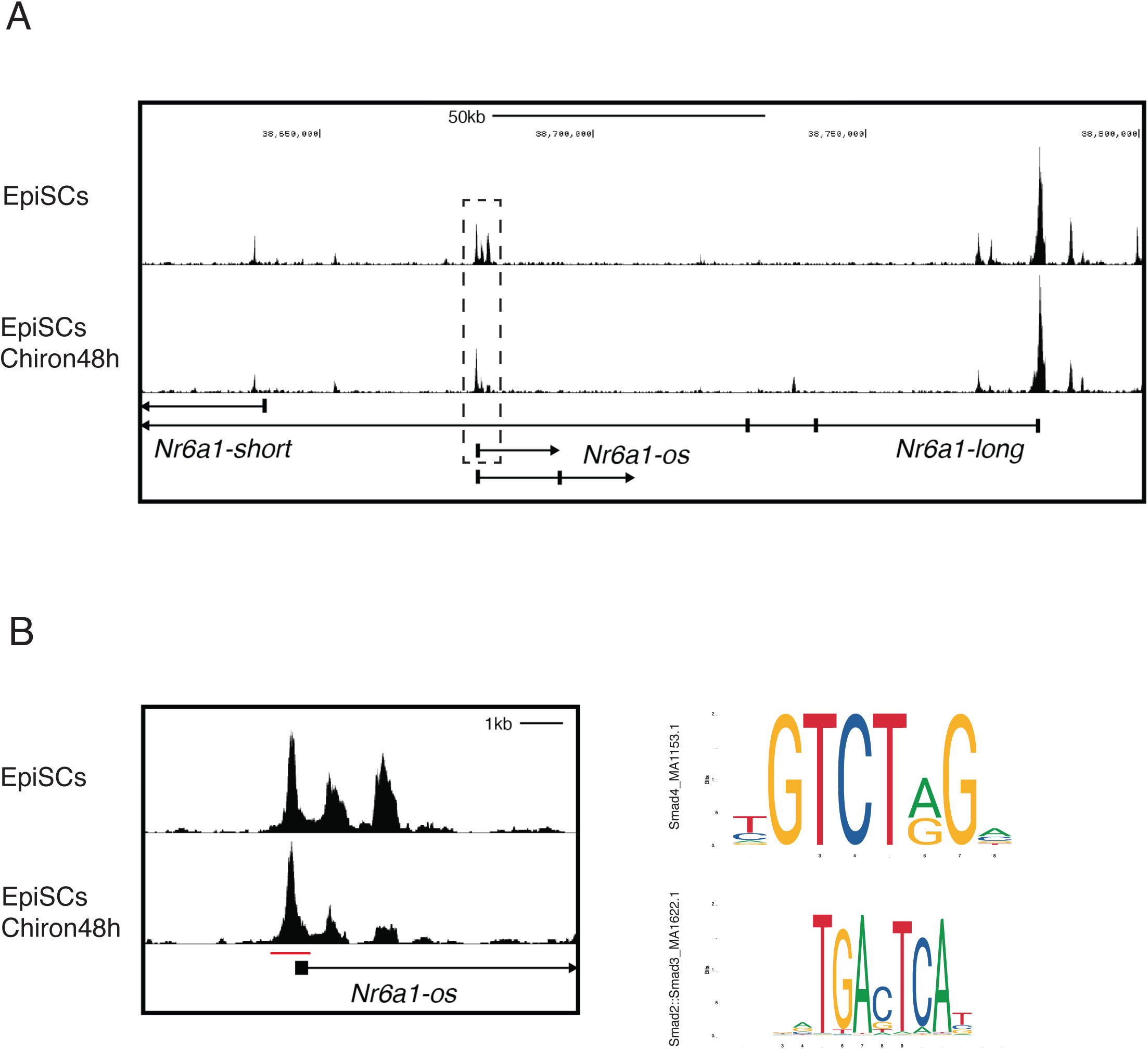
In vitro regulation of *Nr6a1* and *Nr6a1os*. (A) Identification of genomically-accessible regions surrounding the *Nr6a1* locus in epiblast stem cells (EpiSCs) with or without Chiron treatment for 48h. ATAC-seq data derived from (Neijts et al., 2016). (B) The promoter region of *Nr6a1os*, framed in Figure S7A. The red line indicates the genomic region used for transcription factor binding profiles using the JASPAR database (Fornes et al., 2020), revealing multiple predicted Smad binding sites.

**Figure S8.**
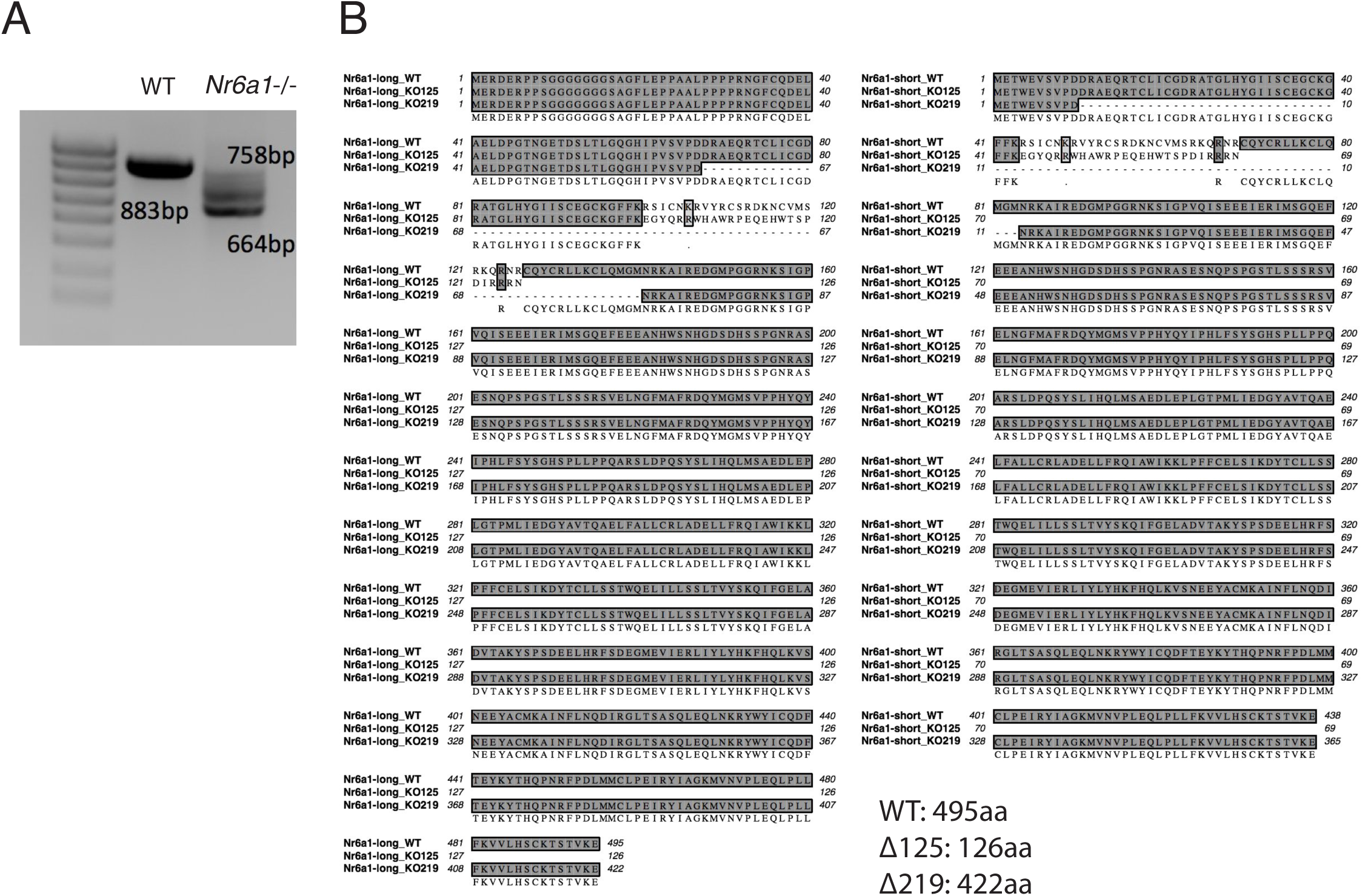
Genomic deletion of *Nr6a1^-/-^* in mouse embryonic stem cells. (A) PCR amplification using primers external to the PAM site reveal successful genomic deletion in one Nr6a1^-/-^ ESC line compared to the expected wildtype product of 883bp. (B) The alignments of wildtype *Nr6a1* with 11219 and 11125 *Nr6a1*.

**Figure S9.**
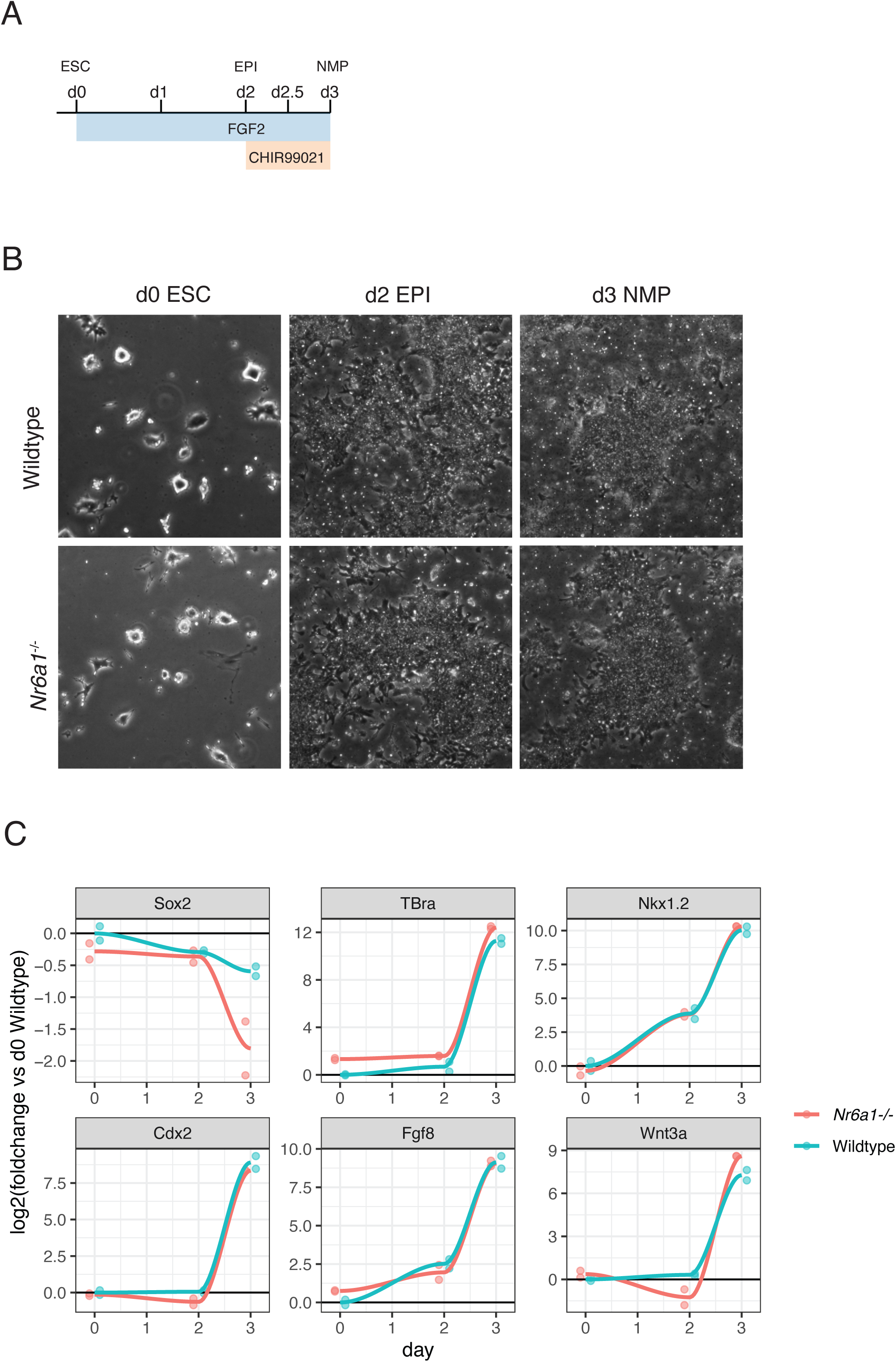
Successful *in vitro* generation of NMPs from *Nr6a1*^-/-^ ECSs. (A) Schematic of *in vitro* differentiation conditions that apply to panels B and C. Embryonic Stem Cell (ESC); Epiblast-like cell (EPI); neuromesodermal-like cell (NMP). (B) At a morphological level, *Nr6a1*^-/-^ cells were indistinguishable from wildtype over the course of *in vitro* differentiation. (C) At a molecular level, *Nr6a1*^-/-^ cells were able to generate NMP-like cells with near-identical kinetics to wildtype, as assessed by qPCR analysis of known NMP marker genes.

**Figure S10.**
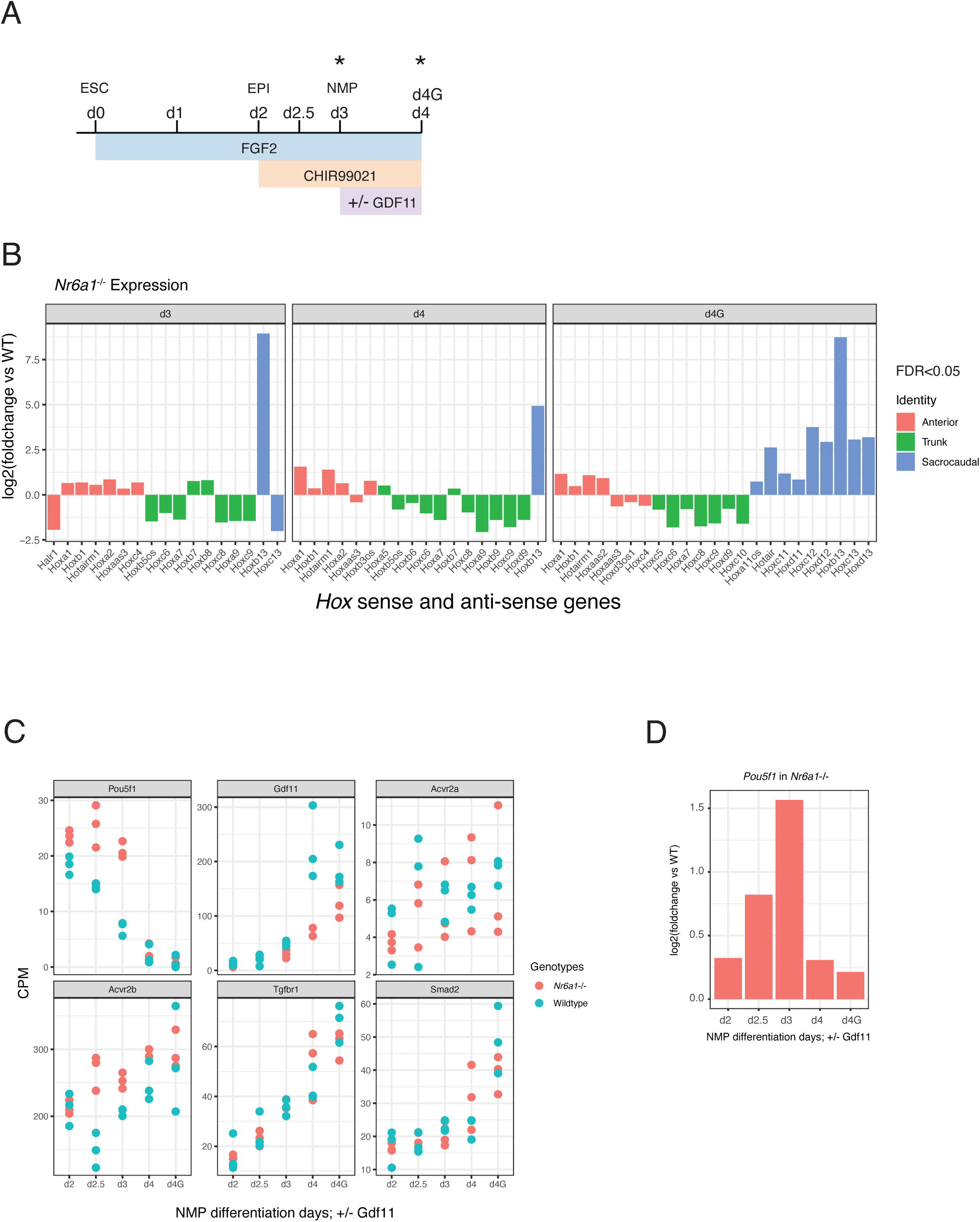
Opposing regulation of trunk vs. caudal *Hox* genes by Nr6a1. (A) Schematic of *in vitro* differentiation conditions that apply to panels B-D. ESC=embryonic stem cell; EPI=epiblast-like cell; NMP=neuromesodermal-like cell. (B) RNAseq analysis of *Nr6a1*^-/-^ in vitro NMPs revealed opposing regulation of trunk vs posterior *Hox* genes by Nr6a1. Results are presented as a log2-transformed fold change in *Nr6a1*^-/-^ cells relative to wildtype, n=3/genotype. Only those *Hox* genes with a false discovery rate (FDR) < 0.05 are displayed, and are colour-coded based on the axial region where the Hox protein functions. (C) RNAseq analysis of *Oct4*/*Pou5f1* and various components of the Gdf11 signaling pathway in *Nr6a1^-/-^* (red) and wild type (blue) cells. Data represented as counts per million (CPM). (D) RNAseq analysis of *Oct4*/*Pou5f1* shown in (C), presented as a log2-transformed fold change in *Nr6a1*^-/-^ samples relative to wildtype. A significant upregulation of *Oct4* in *Nr6a1*^-/-^ cells relative to wildtype was observed at Day(d)2.5 and d3 (FDR < 0.05).

